# TMAO, a seafood-derived molecule, produces diuresis and reduces mortality in heart failure rats

**DOI:** 10.1101/2020.03.18.986869

**Authors:** Marta Gawrys-Kopczynska, Marek Konop, Klaudia Maksymiuk, Katarzyna Kraszewska, Ladislav Derzsi, Krzysztof Sozański, Robert Holyst, Marta Pilz, Emilia Samborowska, Leszek Dobrowolski, Kinga Jaworska, Izabella Mogilnicka, Marcin Ufnal

**Affiliations:** Department of Experimental Physiology and Pathophysiology, Laboratory of the Centre for Preclinical Research, Medical University of Warsaw, Warsaw, Poland; Department of Soft Condensed Matter, Institute of Physical Chemistry, Polish Academy of Sciences, Warsaw, Poland; Mass Spectrometry Laboratory, Institute of Biochemistry and Biophysics, Polish Academy of Sciences, Warsaw, Poland; Department of Renal and Body Fluid Physiology, M. Mossakowski Medical Research Centre, Polish Academy of Sciences, Warsaw, Poland

**Keywords:** Heart failure, SHHF, trimethylamine, trimethylamine N-oxide

## Abstract

**Background:** There is an ongoing debate whether trimethylamine-oxide (TMAO), a molecule present in seafood and a derivate of microbiota metabolism, is beneficial or harmful for the circulatory system. Interestingly, deep-water animals accumulate TMAO that protects proteins such as lactate dehydrogenase (LDH) against high hydrostatic pressure. We hypothesized that TMAO may benefit the circulatory system by protecting cardiac LDH exposed to hydrostatic stress (HS) produced by contracting heart.

**Methods and Results:** Male, 6-week-old, Sprague-Dawley (SD, n=40) and Spontaneously-Hypertensive-Heart-Failure (SHHF n=18) rats were divided into either Water or TMAO oral treatment. After 56 weeks, half of Water and TMAO SD rats were given isoprenaline (ISO) to produce catecholamine stress. In vitro, LDH with or without TMAO was exposed to HS (changes in pressure 0-250mmHg x 280min^−1^) and was evaluated using fluorescence correlation spectroscopy. After 58 weeks of the treatment survival was 100% in SD-Water, SD-TMAO, ISO-TMAO and 90% in ISO-Water. In SHHF-Water survival was 66% vs 100% in SHHF-TMAO. In general, TMAO-treated rats showed higher diuresis and natriuresis. In comparison to SHHF-Water, SHHF-TMAO showed significantly lower diastolic arterial blood pressure, plasma NT-proBNP and expression of angiotensinogen and AT1 receptors in the heart. In separate experiments, intravenous TMAO but not vehicle or urea significantly increased diuresis in SD. In vitro, exposure of LDH to HS with or without TMAO did not affect the protein structure.

**Conclusions:** TMAO reduces mortality in SHHF rats that is associated with diuretic, natriuretic and hypotensive effects. HS produced by the contracting heart is neutral for cardiac LDH structure.

## INTRODUCTION

Plasma TMAO originates from the liver oxidation of trimethylamine (TMA), a product of gut bacteria metabolism of l-carnitine and choline [29, 36]. Furthermore, a direct source of TMAO in humans is TMAO-rich seafood [3, 32].

Several studies showed that increased plasma TMAO levels are associated with an increased risk of adverse cardiovascular events [25–27]. However, other studies have not confirmed this relationship [14, 24, 34]. Furthermore, there are contradictory experimental data regarding the effect of TMAO on the circulatory system [2, 4, 7, 16, 21].

Interestingly, studies in marine animals and biophysical experiments showed protective effect of TMAO. Marine animals living in deep water, and therefore exposed to hydrostatic stress (HS), accumulate TMAO [13, 20, 32, 33, 35]. It has been postulated that the beneficial effect of TMAO depends on stabilization of proteins exposed to HS. For example, TMAO has been found to stabilize teleost and mammalian lactate dehydrogenase (LDH) [33].

We hypothesized that TMAO may benefit the circulatory system by protecting cardiac LDH that is exposed to HS produced by diastole-systole change in hydrostatic pressure in the contracting heart [28]. Notably, in hypertensive heart or in a normotensive heart exposed to catecholamine stress, a diastole-systole change may exceed 220 mmHg, with a frequency of up to 200/min in humans, and higher in small animals. This produces environment in which hydrostatic pressure changes 100 000 times per 24 hours.

Here, we investigated the effect of a chronic,12-months-long oral administration of TMAO in healthy Sprague-Dawley rats (SD), in SD exposed to catecholamine stress and in animal model of heart failure (HF) with reduced ejection fraction i.e. Spontaneously-Hypertensive-Heart-Failure rats (SHHF).

In vitro, we investigated the effect of HS with and without TMAO on the protein structure of cardiac LDH, a complex tetramer protein playing an essential role in cardiac cells metabolism. To evaluate the effect of HS mimicking that produced by the contracting heart we have constructed a novel experimental system which consists of microfluidics chambers with piezoelectric valves and pressure controllers.

## METHODS

### Animals

The study was performed according to Directive 2010/63 EU on the protection of animals used for scientific purposes and approved by the Local Bioethical Committee in Warsaw (permission:100/2016 and 098/2019). 4-5-week-old, male, lean Spontaneously Hypertensive Heart Failure (SHHF/MccGmiCrl-Leprcp/Crl) rats were purchased from Charles River Laboratories (USA). Age-matched Sprague-Dawley (SD) rats were obtained from the Central Laboratory for Experimental Animals, Medical University of Warsaw, Poland.

### Study protocol

Six-week-old SHHF (n=18) and SD (n=40) were randomly assigned to either Water group (rats drinking tap water) or TMAO group (rats drinking TMAO solution in tap water, TMAO-abcr GmbH - Karlsruhe, Germany, 333 mg/l). The dose of TMAO have been selected in order to increase plasma TMAO concentration by 3-5 times (to mimic possible physiological concentrations) and to avoid suprapharmacological effects of TMAO, based on our previous study [7].

Rats were housed in groups of 2-3 animals, in polypropylene cages with environmental enrichment, 12hrs light/12hrs dark cycle, temperature 22-23°C, humidity 45-55%, fed standard laboratory diet (0.19 % Na, Labofeed B standard, Kcynia, Poland) and water ad libitum.

SHHF-TMAO (n=9), SHFF-Water (n=9), SD-TMAO (n=10), SD-Water (n=10) were not subjected to any interventions except of standard animal care until the age of 58 weeks. At the age of 56 weeks ISO-Water (n=10) and ISO-TMAO (n=10) series were given (s.c.) isoprenaline at a dose of 100 mg/kg b.w. (isoprenaline hydrochloride, Sigma-Aldrich, Saint Louis, MO, USA) to produce catecholamine stress as previously described by others [19]. The experimental protocol is depicted in Fig. 1.

**Fig. 1.**
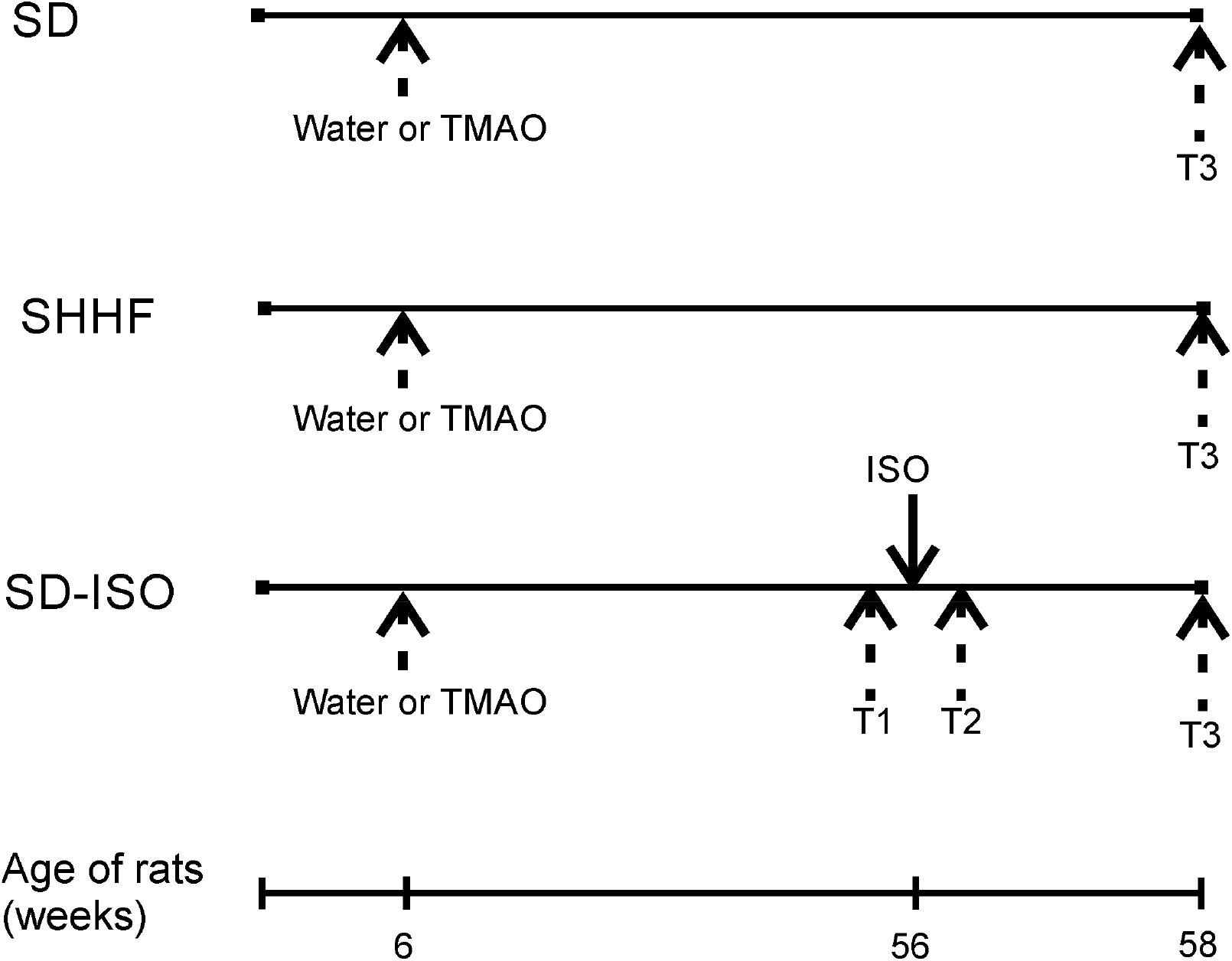
Schematic illustration of experimental series. 6-week-old rats started drinking either Water (Water series) or TMAO water solution (TMAO series). SHHF - Spontaneously Hypertensive Heart Failure (SHHF/MccGmiCrl-Leprcp/Crl) SHHF, SD – Sprague-Dawley rats, SD-ISO - SD rats treated with ISO at the age of 56 week. ISO - administration of isoprenaline at a dose of 100 mg/kg s.c. T1 - metabolic and echocardiographic measurements, T2 - echocardiographic measurements, T3 - metabolic, echocardiographic and direct hemodynamic measurements.

### Experimental protocol in SD and SHHF

58-week-old rats were maintained in metabolism cages for 2 days to evaluate 24hr water and food balance and to collect urine for analysis. Next day, the rats underwent echocardiography using a Samsung HM70. Ultrasound system equipped with a linear probe 5-13. MHz After the echo examination the rats were anaesthetized with urethane (1.5 g/kg b.w. i.p., Sigma-Aldrich) and left femoral artery was cannulated with polyurethane catheter for arterial blood pressure (ABP) recordings. The recordings were started 40 min after the induction of anesthesia and 15 min after inserting the arterial catheter. After 10 min of ABP recordings, Millar Mikro-Tip SPR-320 (2F) pressure catheter was inserted via the right common carotid artery and simultaneous left ventricle pressure (LVP) and ABP recordings were performed. The catheter was connected to Millar Transducer PCU-2000 Dual Channel Pressure Control Unit (Millar, USA) and Biopac MP 150 (Biopac Systems, USA). After hemodynamic recordings, blood from the right ventricle of the heart was taken and rats were killed by decapitation. The heart, the lungs and the kidneys were harvested for histological and molecular biology analysis.

### Experimental protocol in SD-ISO

56-week-old rats were put into metabolic cages for 2 days to evaluate 24hr water and food balance and to collect urine for analysis. Echocardiography was performed as describe above. Next day rats were given isoprenaline (100 mg/kg, s.c.). 24hrs after the administration of ISO the echocardiography was repeated. Eight days after the ISO-treatment 24hr food and water balance were evaluated and echocardiography was performed. Afterwards the rats were anaesthetized with urethane (1.5 g/kg b.w. i.p., Sigma-Aldrich, Poland) and were subjected to hemodynamic measurements including ABP and LVP recordings as described above for SHHF and SD rats.

### TMAO and general biochemistry evaluation

Plasma and urine concentrations of TMAO were measured using Waters Acquity Ultra Performance Liquid Chromatograph coupled with Waters TQ-S triple-quadrupole mass spectrometer. The mass spectrometer operated in the multiple-reaction monitoring (MRM) - positive electrospray ionization (ESI) mode, as previously described [10].

Serum and urine sodium, potassium, creatinine and urea were analyzed using Cobas 6000 analyzer (Roche Diagnostics, Indianapolis, USA).

### ELISA test

The following ELISA kits were used for the evaluation: NT-proBNP (FineTest, cat. no. ER0309), aldosterone (Cayman Chemicals, cat. no. 501090), vasopressin (Biorbyt, cat. no. orb410987), angiotensin II (FineTest, cat. no. ER1637). All procedures were carried out according to the standard protocol supplied with the ELISA Kit. The absorbance intensity was measured at 450 nm with the Multiskan Microplate Reader (Thermo Fisher Scientific). All experiments were performed in doubles.

### Histopathological evaluation

Tissues sections were fixed in 10% buffered formalin, dehydrated using graded ethanol and xylene baths and embedded in paraffin wax. Sections of 3–4 μm were stained with haematoxylin and eosin (HE) and van Gieson stain (for connective tissue fibres). General histopathological examination was evaluated at magnification of 10x, 40x and 100x (objective lens) and 10x (eyepiece) and photographic documentation was made. Morphometric measurements were performed at magnification of 40x (objective lens).

### Molecular Biological Procedures

Heart and kidney samples were collected from rats under urethane anesthesia and frozen in −80°C. Next, samples were homogenized on BeadBug™ microtube homogenizer (Benchmark Scientific, Inc.). Total RNA was isolated from samples according to TRI Reagent® protocol. cDNA was transcribed from RNA samples according to iScript™ Reverse Transcription Supermix #1708841 protocol (Bio-Rad). The qPCR mixes were prepared according to the Bio-Rad SsoAdvanced™ Universal SYBR® Green Supermix protocol #1725271. Amplifications were performed on Bio-Rad CFX Connect Real-Time System under standardized conditions using commercial assays. Data were analyzed by the CFX Manager 3.0 software. Genes investigated in this study were: angiotensinogen (ATG, qRnoCED0051666), angiotensin II receptor type 1a (AT1a, qRnoCID0052626), angiotensin II receptor type 1b (AT1b, qRnoCED0005729), angiotensin II receptor type 2 (AT2, qRnoCED0007551), transforming growth factor-beta (TGF-b, qRnoCED0007638), renin (Rn, qRnoCID0008721), metalloproteinase inhibitor 2 (TIMP2, qRnoCID0001559). Beta-actin was used as housekeeping gene (Actbl2, qRnoCED0018219).

### The effect of TMAO on diuresis, acute experiments

#### Surgical preparations

Male SD rats were anaesthetized with urethane 1.5 g/kg b.w. i.p., which provided stable anesthesia for at least 4 hours. The jugular vein was cannulated for fluid infusions, and the carotid artery for ABP measurement with Biopac MP 150 (Biopac Systems, USA). The bladder was exposed from an abdominal incision and was cannulated for timed urine collection. After the end of surgery, 20-30 min was allowed for stabilization. During this time, 0.9% saline was infused intravenously at a rate of 5 ml/kg/h. After experiments the rats were killed and the both kidneys were excised and weigh.

#### Experimental procedures and measurements

At the end of stabilization period, three or four 10-min urine collections were made to determine baseline water, sodium and total solute excretion rates in each experimental group. After stabilization of urine flow, TMAO (n=8) was infused first as a priming dose of 2.8 mmol/kg b.w. in 5 mL / kg b.w. of 0.9% saline / 5 min, followed by an infusion delivering 2.8 mmol/kg b.w. of TMAO in 5 mL/kg b.w of saline / 60 min. Beginning from the start of the priming injection, five 10 min urine collections were made during the infusion of TMAO. This basic protocol was applied in the two following protocols where TMAO was replaced by its solvent (0.9% NaCl) or saline solution of urea (2.8 mmol/kg).

i. Effects of drug solvent infusion (n = 5). This group served as a control for the equivalent volume administration of fluid bolus.
ii. Effects of hypertonic fluid infusion of urea (n=6). This group served as a control for equivalent volume administration of equivalent hypertonic fluid. The urea solution was equimolar with the solution of TMAO.

#### Analytical procedures and calculations

Urine volume was determined gravimetrically. Urinary osmolality (Uosm) were measured with the cryoscopic Osmomat 030 osmometer (Gonotec, Berlin, Germany). Urine sodium (UNa) and potassium (UK) concentration were measured by a flame photometer (BWB-XP, BWB Technologies Ltd, Newbury, UK). Urine flow (V), the excretion of total solutes (UosmV), sodium (UNaV) and potassium (UKV) were calculated from the usual formulas and standardized to g kidney weight (UXV/g KW).

### Oscillatory-pressure controller and fluorescence correlation spectroscopy

We evaluated the effect of TMAO on cardiac LDH (Merck, Poland) exposed to pressure oscillations and increased temperature. The pressure oscillations were generated in a custom-built system. In order to mimic the conditions in the heart the pressure changed from 0 to 180–250 mmHg (or to higher values) at oscillation rate of 280 min-1. In general, the setup consisted of two main parts: i) a custom-built oscillatory pressure controller with solenoid micro valves to control the inner pressure and “pulse” frequency and ii) sample chamber (Fig. 2). We designed and constructed 3 different samples chambers (Fig. 3,4,5), which allowed us to expose the samples to pressure oscillation in different ways. Details on the construction and parameters of the pressure oscillation systems can be found in the Supplemental Material (Fig. S1-S6).

**Fig. 2.**
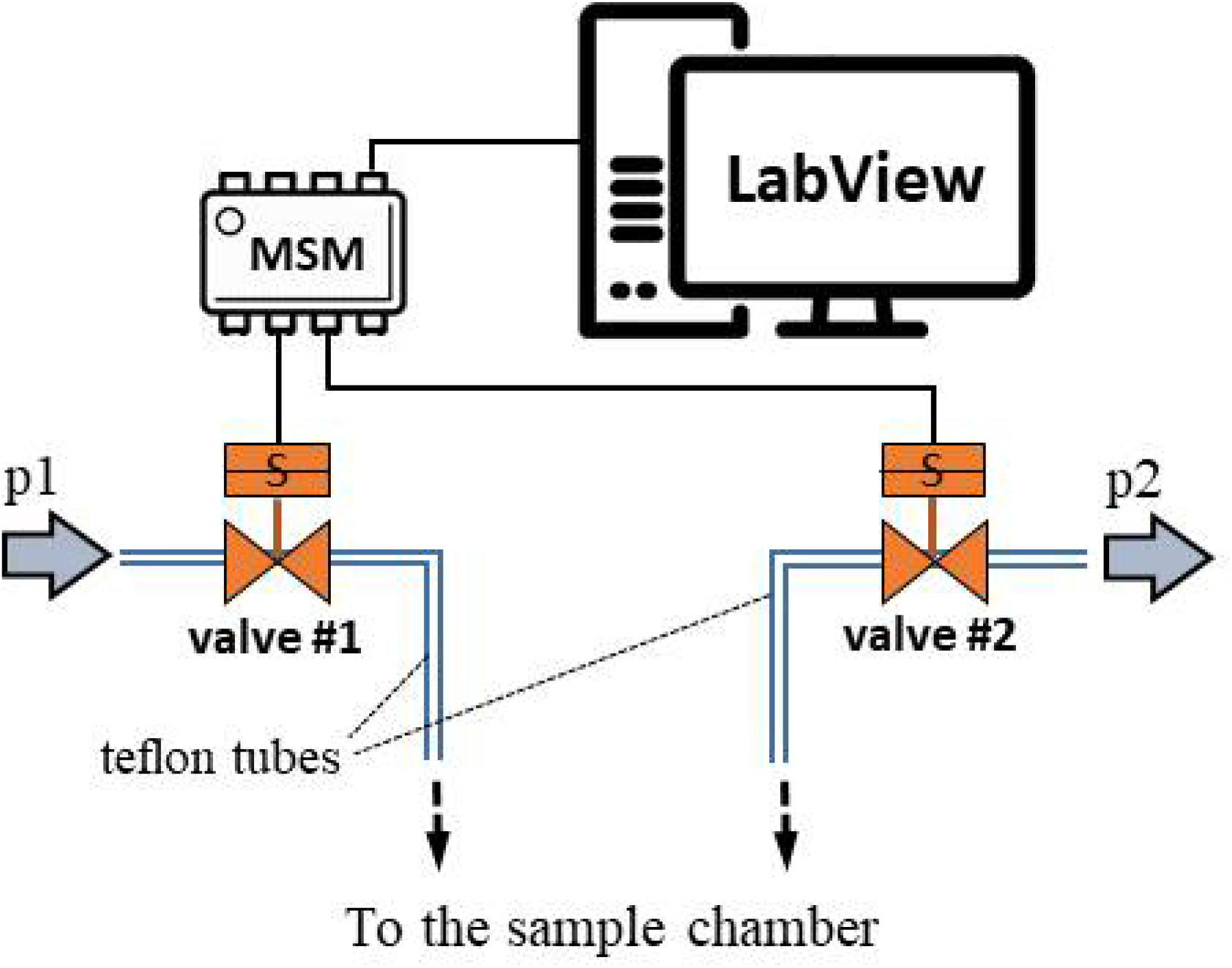
Schematic illustration of the oscillatory pressure controller.

**Fig. 3.**
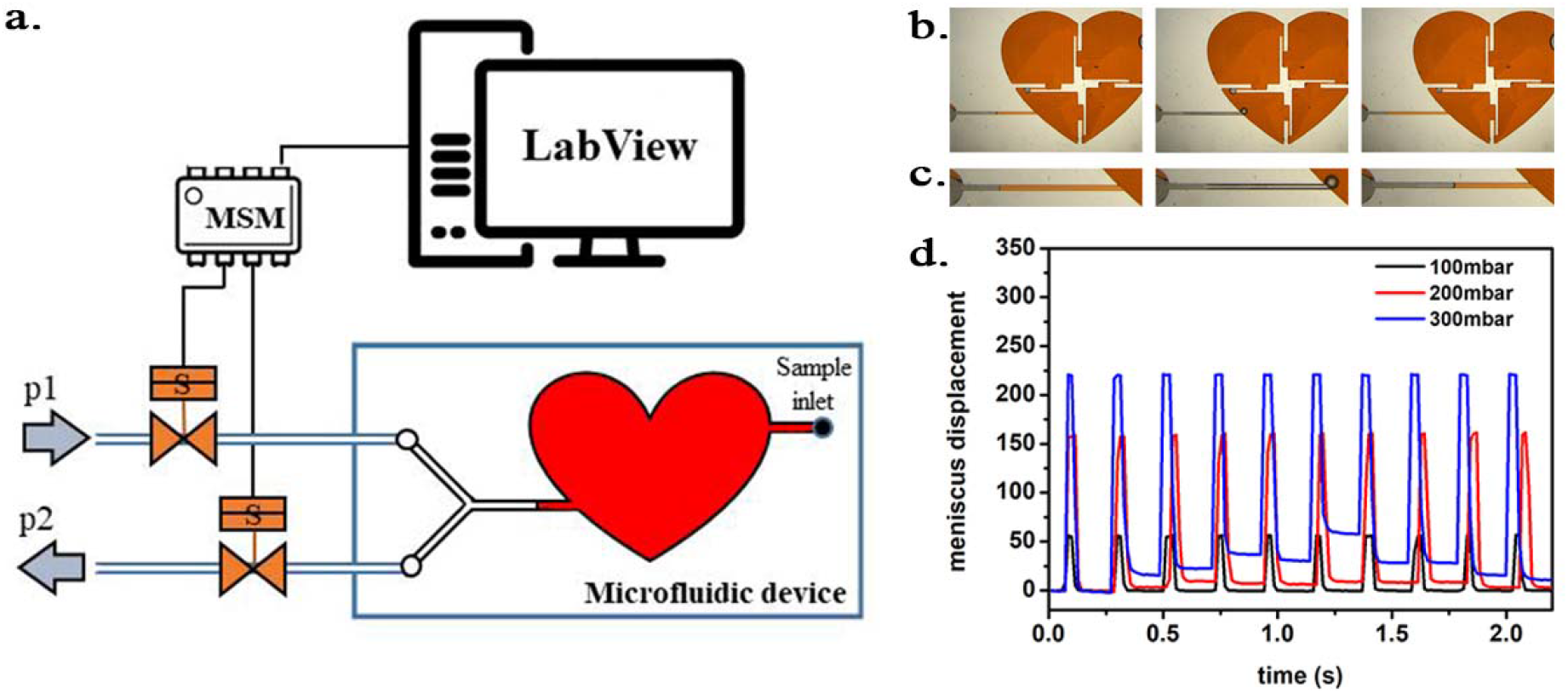
a) Scheme of the experimental setup with the heart-shaped PDMS microfluidic device. b) time-sequence snapshots of the device in operations. From left to right: the device is filled with liquid and is at rest (for better visualisation we used here red-dyed water instead of the transparent protein solution). The microchannel connecting the pressure system to the microfluidic chamber is half-way filled. Applying high pressure (valve 1 open) pushes the liquid meniscus toward the chamber. Closing valve #1 and opening valve #2 (low pressure) the liquid meniscus pulls back (even further than its original position). c) Close-up of the moving liquid meniscus, d) oscillation profile constructed from the position of the liquid meniscus in the microchannel as a function of time for various pressure differences *Δp=p1-p2*.

**Fig. 4.**
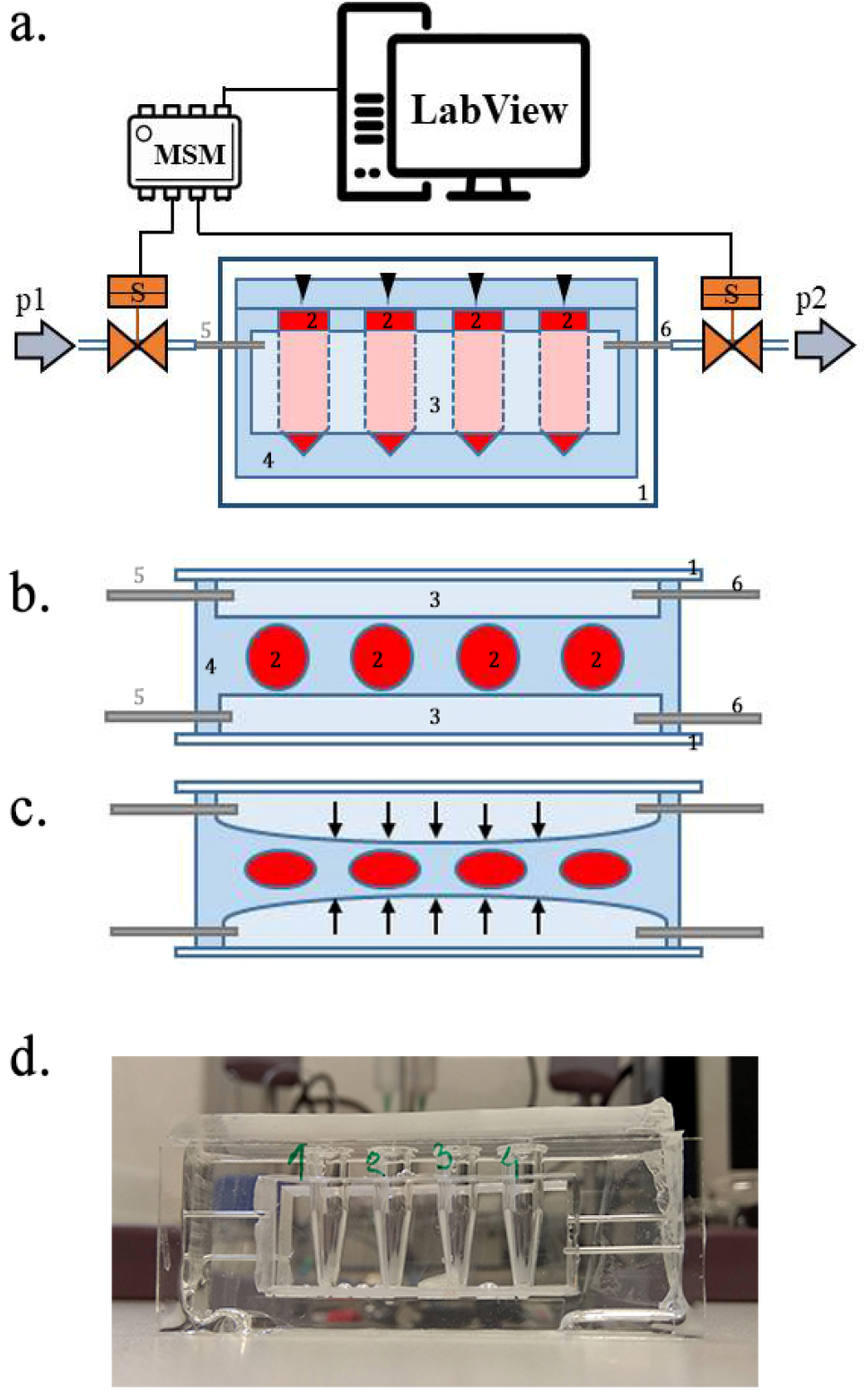
a) Scheme of the set-up with the pressure oscillator connected to the Eppendorf chip with side view and b) top view of the “Eppendorf chip”. Numbers indicating the glass slide (1), the sample chambers (2), the cuboid cavity (3), the PDMS and the inlet (5) and outlet (6) steel capillaries, respectively. b-c) Schematic representation of the chip’s operation upon applied pressure: the cavity (3) expands towards the sample chambers (2) and squeeze them. d) Photo of the constructed device prior to filling and connection to the pressure controller.

**Fig. 5.**
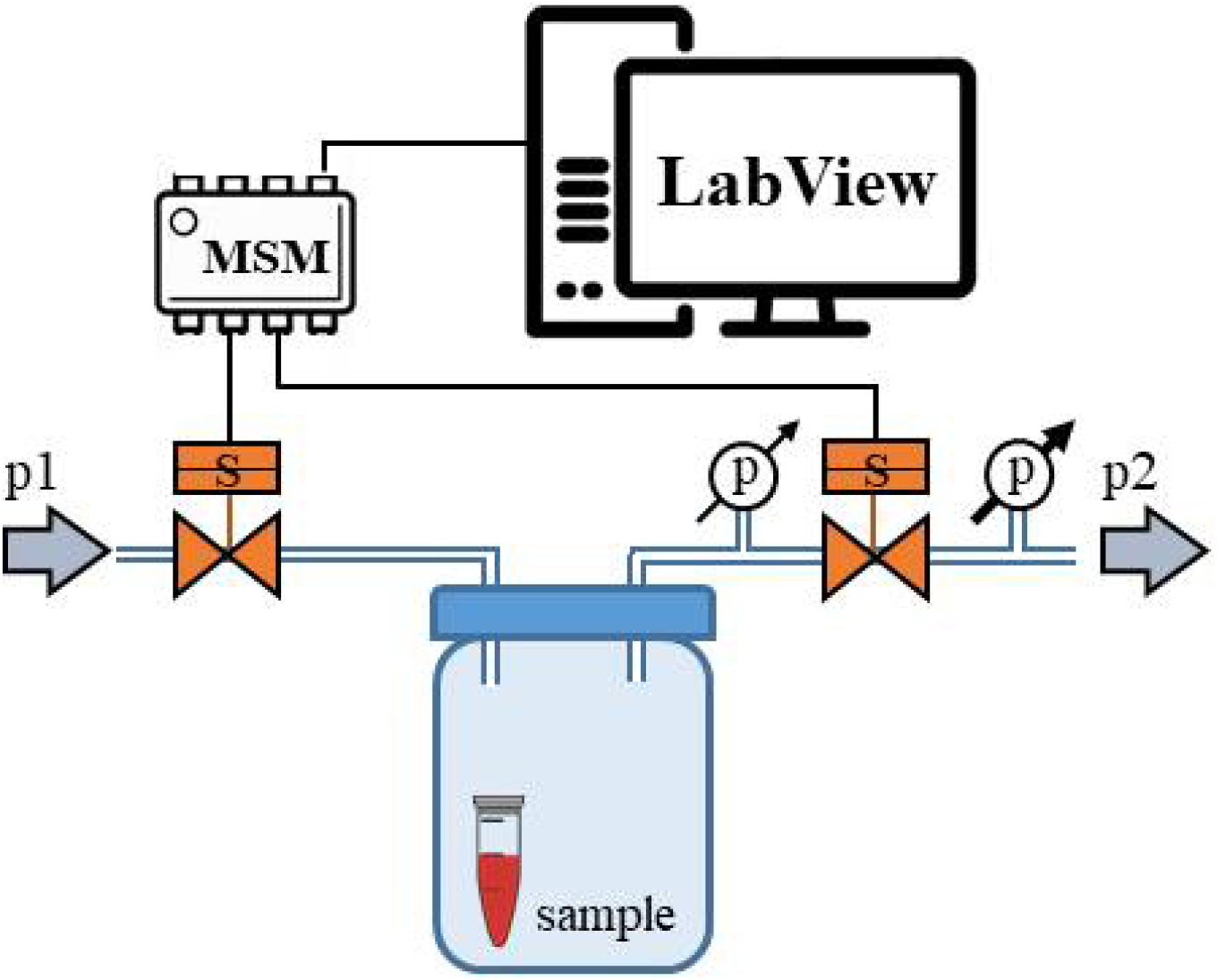
Schematic representation of the “pressure bottle” system. Two manometers were attached to the system and used to determine the pressure acting on the samples.

In all experiments, proteins at a concentration of around 1 μM were incubated in the pressure oscillation system for 24 hrs at room temperature. In each experimental run, we incubated in parallel proteins suspended in pure phosphate buffer saline (PBS) and 1M solution of TMAO in PBS. As a control for each of the samples, we used the same solution placed in a chamber same as the pressure oscillation chamber and in the same conditions, but under constant, atmospheric pressure. All solutions were then diluted 100-fold in PBS and fluorescence correlation spectroscopy (FCS) measurements were performed.

The effect of hydrodynamic and thermal stress on Atto 488 (ATTO-TEC GmbH) labeled LDH structure was evaluated by means of FCS. Protein diffusion coefficients (*D*) is associated with its hydrodynamic radius (*R*_*h*_) by Stokes-Einstein relation, *D* = *k_B_T*/6πη*R*_*h*_ (where *k*_*B*_ is Boltzmann constant, *T* is the temperature and *η* denotes the viscosity of medium). The alteration in protein structure as dissociation of the quaternary structure results in an increase of the observed diffusion coefficient, while denaturation of tertiary structure causes a decrease of *D* by a factor of 1.5 – 3 [30].

FCS measurements were performed on dedicated FCS system, based on Nikon C1 inverted confocal microscope (Nikon Instruments, Japan) with PlanApo 60x, NA=1.2 water immersion objective. The setup is equipped with Pico Harp 300 system (PicoQuant, Germany). Measurements were performed in a climate chamber (Okolab, Italy) providing temperature control and required humidity at 25.0 ± 0.5°C. Labeled protein was excited by a 485 nm laser, and fluorescence was detected through a 488/LP long-pass filter (Chroma, USA). Data acquisition and analysis was performed in SymPhoTime 64 software (PicoQuant, Germany). The experiments were preceded by establishing the dimension of the confocal volume using rhodamine 110 (Sigma-Aldrich, USA).

### Data analysis and statistics

Diastolic arterial blood pressure (DBP), systolic arterial blood pressure (SBP) and heart rate (HR) were calculated on the arterial blood pressure tracing. Left ventricular end-diastolic pressure (LVEDP), maximal slope of systolic pressure increment (+dP/dt) and diastolic pressure decrement (-dP/dt) were calculated on the left ventricle blood pressure tracing using AcqKnowledge Biopac software (Biopac Systems, Goleta, USA). Differences between the TMAO and Water treatment were evaluated by independent-samples t-test. The log-rank test was used to test the survival differences between TMAO and Water treatment. A value of two-sided P<0.05 was considered significant. Analyses were conducted using Statistica, version 13.3 (Tibco, Palo Alto, CA, USA).

## RESULTS

### SD rats: Water vs TMAO treatment

In general SD rats showed no pathological findings (Table 1, Fig. 6,7 and supplemental Fig. S7).

**Table 1.**
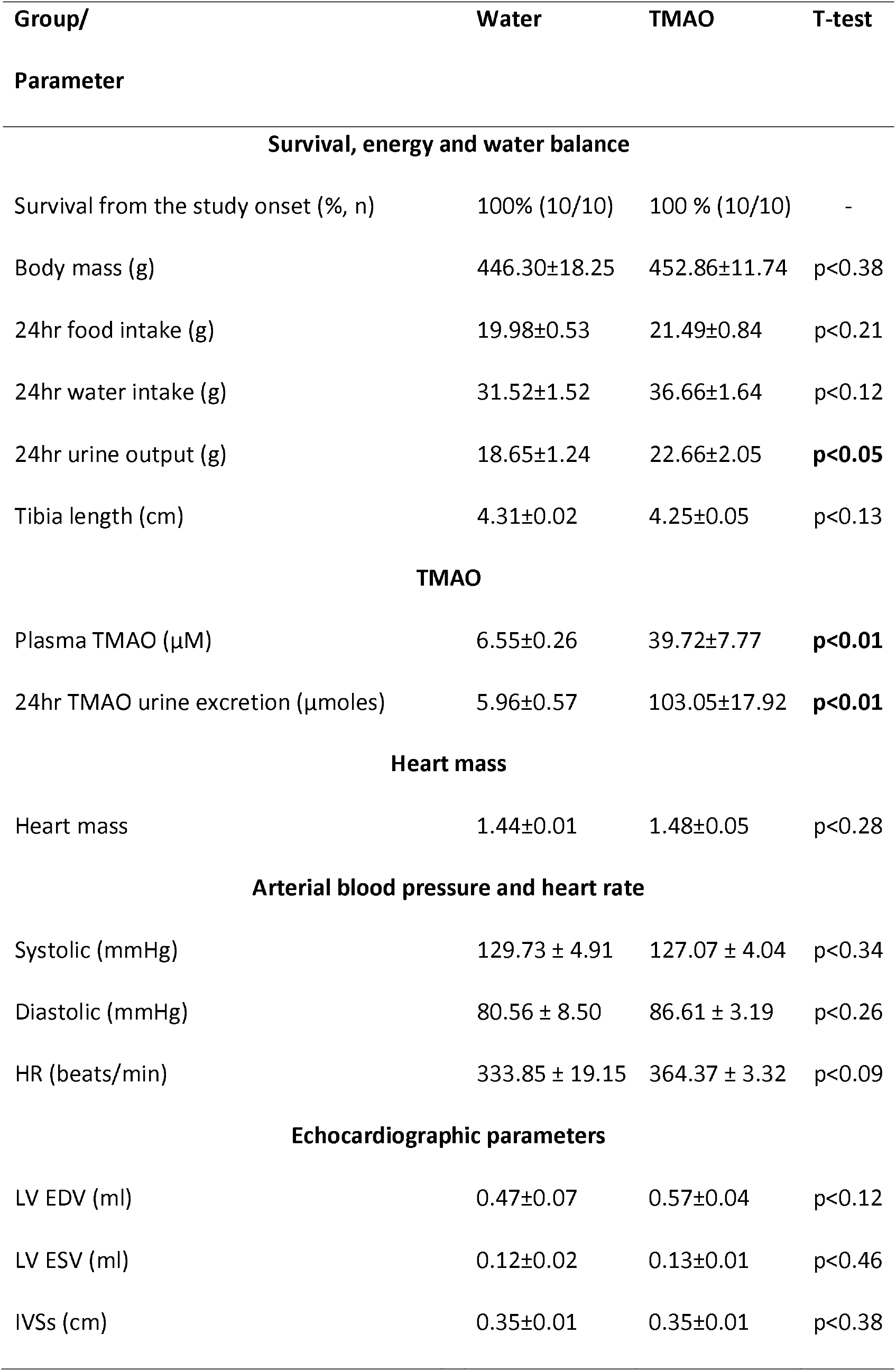
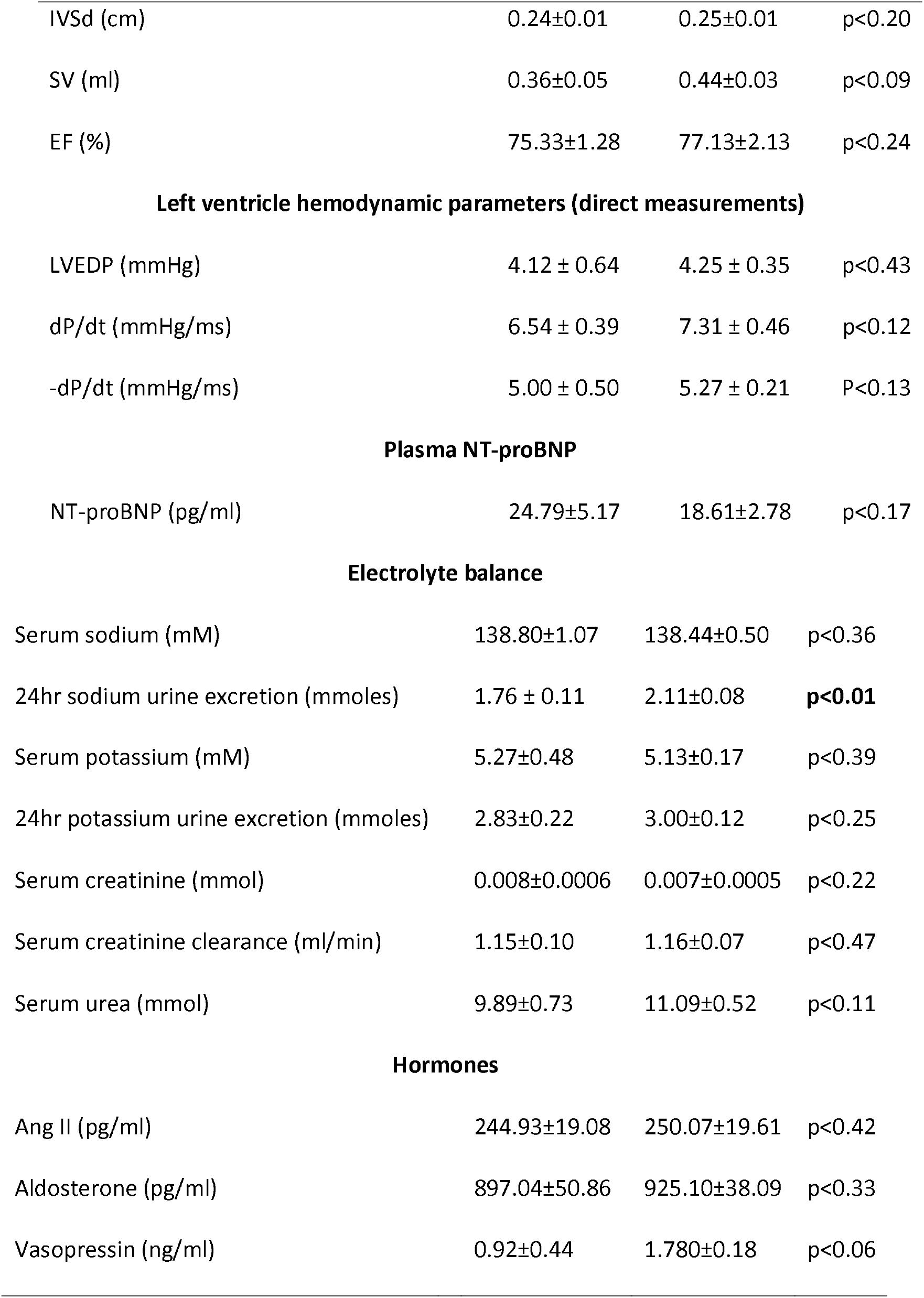
Metabolic, renal and cardiovascular parameters in 58-week-old normotensive Sprague-Dawley rats maintained on either Water or TMAO solution in drinking water. Creatinine clearance calculated as urine creatinine x urine output (ml/min) / plasma creatinine. LVEDV - left ventricle end diastolic volume, LVESV - left ventricle end systolic volume, SV – stroke volume, EF - ejection fraction, IVSs(d) - intraventricular septum diameter during systole and diastole, respectively, LVEDP - pressure in the left ventricle during the end of diastole measured directly with a catheter, +dP/dt - maximal slope of systolic ventricular pressure increment, - dP/dt - maximal slope of diastolic ventricular pressure decrement. Values are means, ± SE. P values by T-test.

| Group/<br>Parameter | Water | TMAO | T-test |
| --- | --- | --- | --- |
| **Survival, energy and water balance** |  |  |  |
| Survival from the study onset (%*, n*) | 100% (10/10) | 100 % (10/10) | - |
| Body mass (g) | 446.30±18.25 | 452.86±11.74 | p<0.38 |
| 24hr food intake (g) | 19.98±0.53 | 21.49±0.84 | p<0.21 |
| 24hr water intake (g) | 31.52±1.52 | 36.66±1.64 | p<0.12 |
| 24hr urine output (g) | 18.65±1.24 | 22.66±2.05 | **p<0.05** |
| Tibia length (cm) | 4.31±0.02 | 4.25±0.05 | p<0.13 |
| **TMAO** |  |  |  |
| Plasma TMAO (μM) | 6.55±0.26 | 39.72±7.77 | **p<0.01** |
| 24hr TMAO urine excretion (μmoles) | 5.96±0.57 | 103.05±17.92 | **p<0.01** |
| **Heart mass** |  |  |  |
| Heart mass | 1.44±0.01 | 1.48±0.05 | p<0.28 |
| **Arterial blood pressure and heart rate** |  |  |  |
| Systolic (mmHg) | 129.73 ± 4.91 | 127.07 ± 4.04 | p<0.34 |
| Diastolic (mmHg) | 80.56 ± 8.50 | 86.61 ± 3.19 | p<0.26 |
| HR (beats/min) | 333.85 ± 19.15 | 364.37 ± 3.32 | p<0.09 |
| **Echocardiographic parameters** |  |  |  |
| LV EDV (ml) | 0.47±0.07 | 0.57±0.04 | p<0.12 |
| LV ESV (ml) | 0.12±0.02 | 0.13±0.01 | p<0.46 |
| IVSs (cm) | 0.35±0.01 | 0.35±0.01 | p<0.38 |
| IVSd (cm) | 0.24±0.01 | 0.25±0.01 | p<0.20 |
| SV (ml) | 0.36±0.05 | 0.44±0.03 | p<0.09 |
| EF (%) | 75.33±1.28 | 77.13±2.13 | p<0.24 |
| Left ventricle hemodynamic parameters (direct measurements) |  |  |  |
| LVEDP (mmHg) | 4.12 ± 0.64 | 4.25 ± 0.35 | p<0.43 |
| dP/dt (mmHg/ms) | 6.54 ± 0.39 | 7.31 ± 0.46 | p<0.12 |
| -dP/dt (mmHg/ms) | 5.00 ± 0.50 | 5.27 ± 0.21 | P<0.13 |
| Plasma NT-proBNP |  |  |  |
| NT-proBNP (pg/ml) | 24.79±5.17 | 18.61±2.78 | p<0.17 |
| Electrolyte balance |  |  |  |
| Serum sodium (mM) | 138.80±1.07 | 138.44±0.50 | p<0.36 |
| 24hr sodium urine excretion (mmoles) | 1.76 ± 0.11 | 2.11±0.08 | **p<0.01** |
| Serum potassium (mM) | 5.27±0.48 | 5.13±0.17 | p<0.39 |
| 24hr potassium urine excretion (mmoles) | 2.83±0.22 | 3.00±0.12 | p<0.25 |
| Serum creatinine (mmol) | 0.008±0.0006 | 0.007±0.0005 | p<0.22 |
| Serum creatinine clearance (ml/min) | 1.15±0.10 | 1.16±0.07 | p<0.47 |
| Serum urea (mmol) | 9.89±0.73 | 11.09±0.52 | p<0.11 |

**Table 1.** Metabolic, renal and cardiovascular parameters in 58-week-old normotensive Sprague-Dawley rats maintained on either Water or TMAO solution in drinking water. Creatinine clearance calculated as urine creatinine x urine output (ml/min) / plasma creatinine.
|  |  |  |  |
| --- | --- | --- | --- |
| Ang II (pg/ml) | 244.93±19.08 | 250.07±19.61 | p<0.42 |
| Aldosterone (pg/ml) | 897.04±50.86 | 925.10±38.09 | p<0.33 |
| Vasopressin (ng/ml) | 0.92±0.44 | 1.780±0.18 | p<0.06 |

**Fig. 6.**
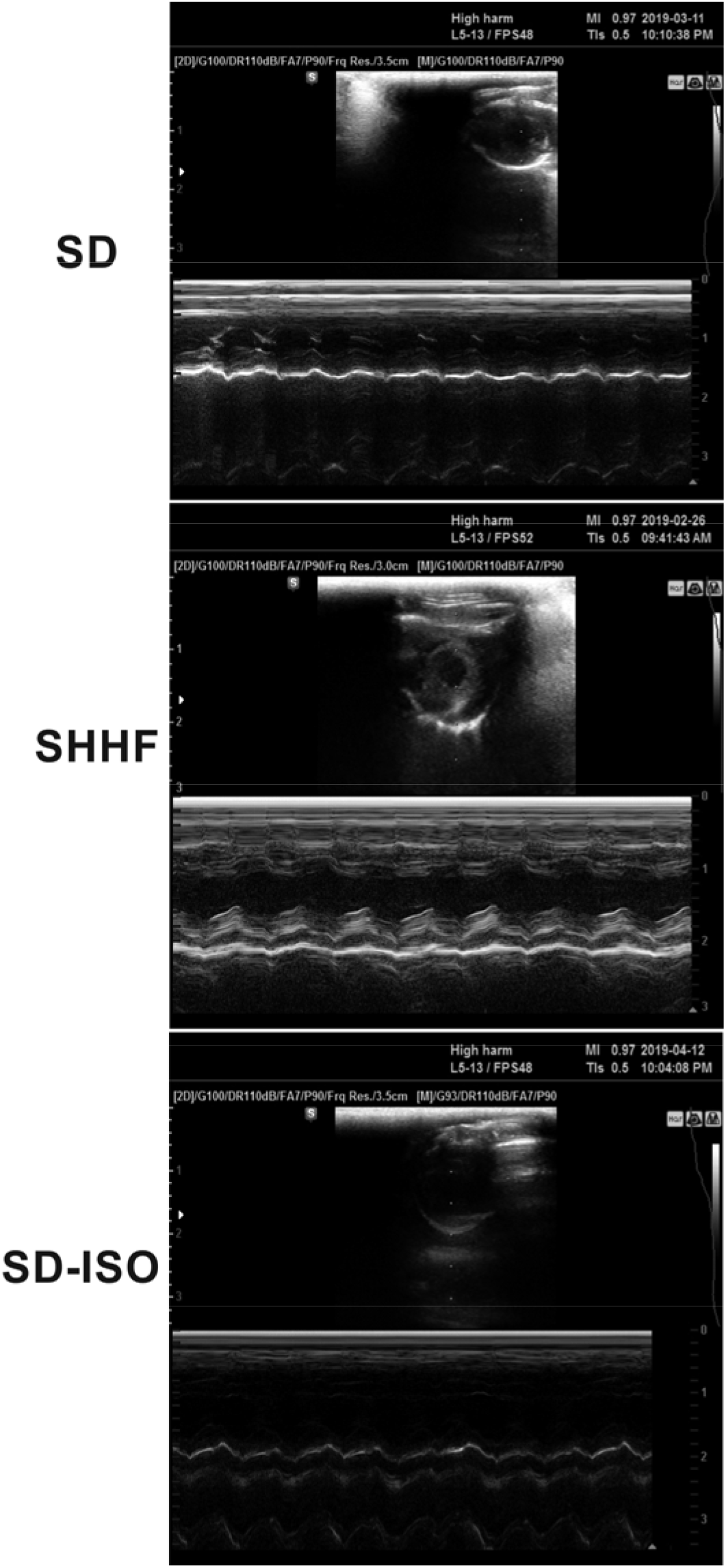
Echocardiography in 58-wk-old rats. SHHF - Spontaneously Hypertensive Heart Failure (SHHF/MccGmiCrl-Leprcp/Crl) SHHF, SD – Sprague-Dawley rats, SD-ISO: SD rats treated with ISO at the age of 56 week. ISO: administration of isoprenaline at a dose of 100 mg/kg s.c. SD: Left ventricular systolic function is normal/preserved. Left and right ventricular diameter is normal. Left ventricular wall thickness is normal. Left atrial diameter is normal. SHHF Septal hypokinesis. Left ventricular free wall is hypertrophic. Endocardium is hyperechogenic. SD-ISO: Septal hypokinesis. Left ventricular end-systolic diameter is increased. Left atrial enlargment.

**Fig. 7.**
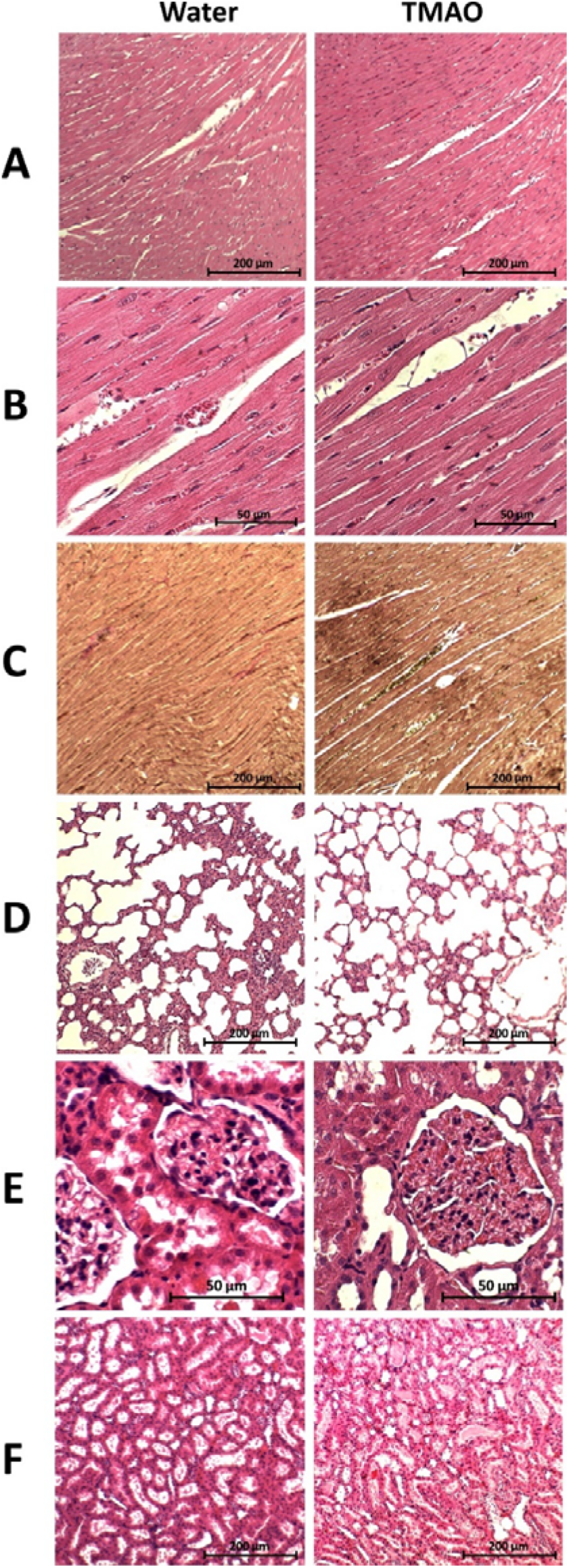
Histopathological picture of heart, lungs and kidneys in Sprague-Dawley rats drinking either Water or TMAO solution. A - myocardium; haematoxylin-eosin staining at magnification x10; B - myocardium; haematoxylin-eosin staining at magnification x40; C - myocardium; van Gieson staining at magnification x10; D - lungs; haematoxylin-eosin staining at magnification x10; E – kidney - renal cortex, renal bodies; haematoxylin-eosin staining at magnification x40; F - kidney - renal medulla; hematoxylin-eosin staining at magnification x10.

#### Survival and water-electrolyte balance

There was no significant difference between Water and TMAO treated rats in survival rate (100% in both groups), body mass and food intake. TMAO-treated rats showed 5-times higher plasma TMAO level than Water-treated rats. The TMAO group showed a significantly higher 24hr urine output and sodium urine excretion. There was no significant difference between TMAO and Water group in plasma Ang II and aldosterone level. However, there was a trend towards higher vasopressin concentration in TMAO rats (p<0.06), (Table 1).

#### Circulatory parameters

There was no significant difference between Water and TMAO group in hemodynamic parameters measured directly and with echocardiography (Table 1).

#### Histopathology

There were no pathological changes in the heart, lungs and kidney in Water and TMAO groups (Fig. 7).

#### Gene expression

TMAO treated rats showed significantly lower expression of AT1 receptors (AT1R) in the heart. In the kidneys TMAO treated rats showed significantly higher expression of renin but lower expression of AT2 receptors and a trend towards lower expression of AT1R (Fig. 8,9).

**Fig. 8.**
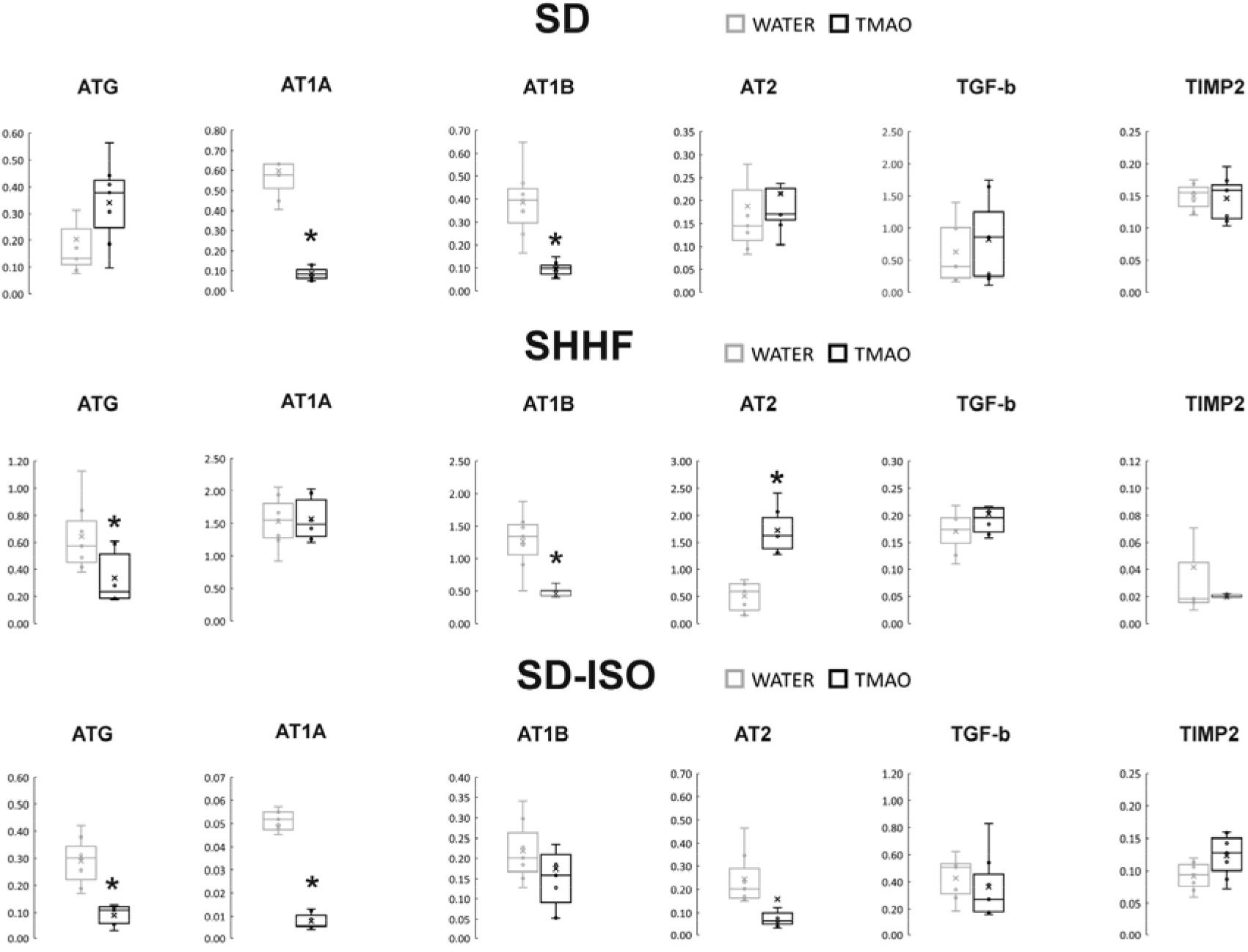
Real-time RT-PCR analysis, heart. Box plot comparing the expression profiles of ATG (angiotensinogen), AT1A (angiotensin II receptor type 1a), AT1B (angiotensin II receptor type 1b), AT2 (angiotensin II receptor type 2), TGF-b (transforming growth factor-beta), TIMP2 (metalloproteinase inhibitor 2) in hearts of SD – Sprague-Dawley rats, SHHF - Spontaneously Hypertensive Heart Failure (SHHF/MccGmiCrl-Leprcp/Crl), SD-ISO - SD rats treated with ISO at the age of 56 week, Water (Water series) or TMAO water solution (TMAO series). × - mean value, * indicates significant difference compared with Water groups. *P < 0.05

**Fig. 9.**
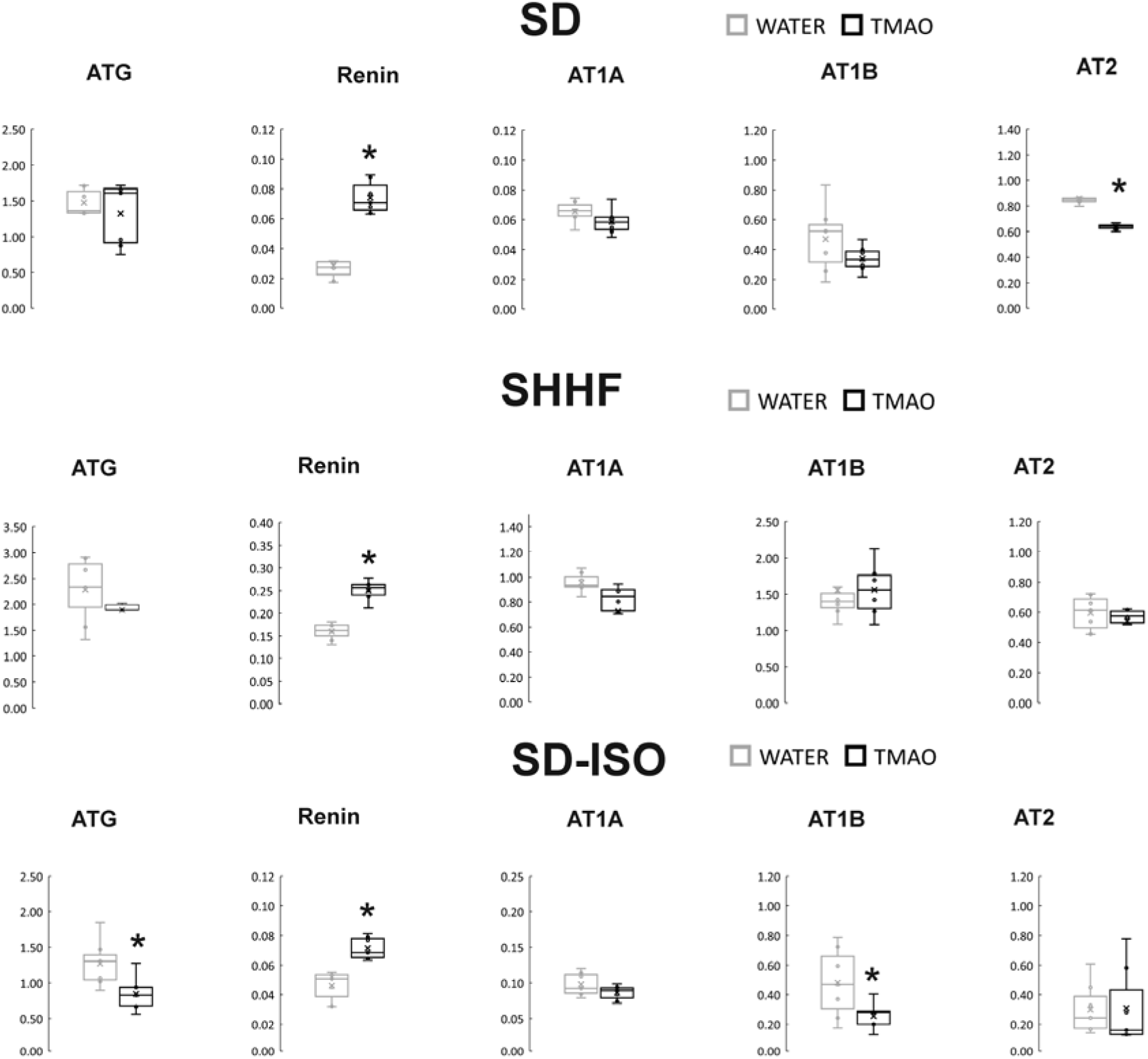
Real-time RT-PCR analysis, kidneys. Box plot comparing the expression profiles of ATG (angiotensinogen), AT1A (angiotensin II receptor type 1a), AT1B (angiotensin II receptor type 1b), AT2 (angiotensin II receptor type 2), renin in kidneys of SD – Sprague-Dawley rats, SHHF - Spontaneously Hypertensive Heart Failure (SHHF/MccGmiCrl-Leprcp/Crl), SD-ISO - SD rats treated with ISO at the age of 56 week, Water (Water series) or TMAO water solution (TMAO series). × - mean value,* indicates significant difference compared with Water groups. *P < 0.05

### SHHF rats: Water vs TMAO treatment

In general, SHHF showed characteristics of hypertrophic cardiomyopathy with compromised systolic function including substantially increased heart mass and plasma NT-proBNP, decreased stroke volume and ejection fraction as well as lung edema (Table 2, Fig. 6). Histopathological picture of SHHF heart corresponds to dilated cardiomyopathy i.e. cardiomyocytes showed a moderate increase in diameter, the enlargement of nucleus and the reduction of cytoplasmic acidity (Fig. 10). In SHHF lungs showed signs of passive hyperemia, with thickening of the interalveolar septa and a weak focal parenchymal edema. Moderate stromal connective tissue hyperplasia was also found. There were no significant pathological changes in the SHHF kidneys (Fig. 10).

**Table 2.** Metabolic, renal and cardiovascular parameters in 58-week-old Spontaneously Hypertensive Heart Failure (SHHF/MccGmiCrl-Leprcp/Crl) rats maintained on either Water or TMAO solution in drinking water. Creatinine clearance calculated as urine creatinine x urine output (ml/min) / plasma creatinine. LVEDV - left ventricle end diastolic volume, LVESV - left ventricle end systolic volume, SV – stroke volume, EF - ejection fraction, IVSs(d), intraventricular septum diameter during systole and diastole, respectively. LVEDP - pressure in the left ventricle during the end of diastole measured directly with a catheter, +dP/dt - maximal slope of systolic ventricular pressure increment, - dP/dt - maximal slope of diastolic ventricular pressure decrement. Values are means, ± SE. p values by T-test, except # - by log-rank test.

| Group/<br>Parameter | Water | TMAO | T-test |
| --- | --- | --- | --- |
| **Survival, Energy and water balance** |  |  |  |
| Survival from the onset of the study (% , n) | 66% (6/9) | 100 % (9/9) | p<0.07 # |
| Body mass (g) | 475.2±7.8 | 476.3±5.1 | p<0.93 |
| 24hr food intake (g) | 23.2±1.5 | 24.2±0.9 | p<0.69 |
| 24hr water intake (ml) | 37.4±3.4 | 41.1±2.8 | p<0.36 |
| 24hr urine output (ml) | 14.8±1.3 | 17.9±1.2 | **p<0.05** |
| Tibia length (cm) | 3.96±0.10 | 3.98±0.40 | p<0.79 |
| **TMAO** |  |  |  |
| Plasma TMAO (μM) | 6.71±1.66 | 20.32±2.95 | **p<0.01** |
| 24hr TMAO urine excretion (μmoles) | 9.97±1.39 | 126.8±11.9 | **p<0.01** |
| **Heart mass** |  |  |  |
| Heart mass | 1.87±0.14 | 1.72±0.12 | p<0.37 |
| **Arterial blood pressure and heart rate** |  |  |  |
| Systolic (mmHg) | 136.2 ± 5.8 | 126.8 ± 5.4 | p<0.26 |
| Diastolic (mmHg) | 98.6 ± 3.4 | 87.6 ± 2.1 | **p<0.01** |
| HR (beats/min) | 314 ± 27 | 302 ± 8 | p<0.59 |
| **Echocardiographic parameters** |  |  |  |
| LV EDV (ml) | 0.37±0.09 | 0.52±0.08 | p<0.21 |
| LV ESV (ml) | 0.14±0.04 | 0.15±0.04 | p<0.82 |
| IVSs (cm) | 0.41±0.01 | 0.35±0.03 | p<0.15 |
| IVSd (cm) | 0.29±0.02 | 0.26±0.03 | p<0.56 |
| SV (ml) | 0.24±0.06 | 0.36±0.05 | p<0.10 |
| EF (%) | 64±4.0 | 71±2.4 | p<0.12 |
| **Left ventricle hemodynamic parameters (direct measurements)** |  |  |  |
| LVEDP (mmHg) | 3.10 ± 0.35 | 3.41 ± 0.33 | p<0.47 |
| dP/dt (mmHg/ms) | 5.87 ± 0.41 | 5.49 ± 0.39 | p<0.69 |
| -dP/dt (mmHg/ms) | 2.34 ± 0.13 | 2.55 ± 0.29 | p<0.62 |
| **Plasma NT-proBNP** |  |  |  |
| NT-proBNP (pg/ml) | 52.26±5.9 | 42.81±3.4 | p<0.24 |
| **Electrolyte balance** |  |  |  |
| Serum sodium (mM) | 142.3±1.65 | 138.9±1.2 | p<0.11 |
| 24hr sodium urine excretion (mmoles) | 1.41 ± 0.12 | 1.72 ± 0.13 | **p<0.01** |
| Serum potassium (mM) | 4.73±0.13 | 4.47±0.11 | p<0.19 |
| 24hr potassium urine excretion (mmoles) | 3.02±0.21 | 2.89±0.19 | p<0.08 |
| Serum creatinine clearance (ml/min) | 0.42±0.07 | 0.53±0.02 | p<0.11 |
| **Hormones** |  |  |  |
| Ang II (pg/ml) | 325.7±19.9 | 276.8±12.8 | **p<0.02** |
| Aldosterone (pg/ml) | 816.8±134.3 | 758.5±51.2 | p<0.64 |
| Vasopressin (ng/ml) | 3.02±0.49 | 3.11±0.42 | p<0.89 |

**Fig. 10.**
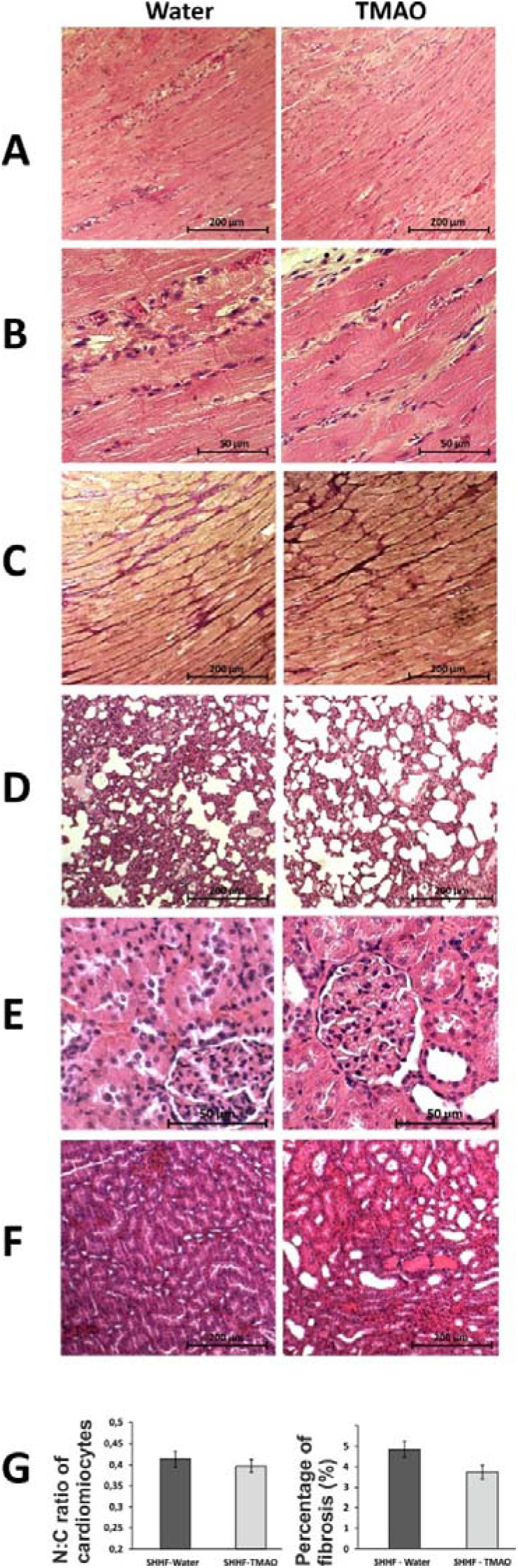
Histopathological picture of heart, lungs and kidneys in Spontaneously Hypertensive Heart Failure (SHHF/MccGmiCrl-Leprcp/Crl) drinking either Water or TMAO solution. A - myocardium; hematoxylin-eosin staining at magnification x10; B - myocardium; hematoxylin-eosin staining at magnificarion x40; C - myocardium; van Gieson staining at magnification x10; D - lungs; hematoxylin-eosin staining at magnification x10; E – kidney - renal cortex, renal bodies; hematoxylin-eosin staining at magnification x40; F – kidney - renal medulla; hematoxylin-eosin staining at magnification x10; G – Percentage of myocardial fibrosis [%], N:C ratio of cardiomyocytes.

#### Survival and water-electrolyte balance

All SHHF-TMAO rats (n=9) survived from the beginning of the experiment till the age of 58-weeks, i.e. the time of anesthesia before echocardiography. In contrast, three out of nine (33%) SHHF-controls were euthanized or died spontaneously at the age of 52, 56 and 57-weeks due to hemiparalysis (ischemic stroke), dyspnoe (post-mortem lung edema) and spontaneous death (post-mortem lung edema), respectively. The log-rank test showed a trend p=0.0651 toward higher survival in SHHF-TMAO than in SHHF-Water (Fig. 11). One SHHF-TMAO died during anesthesia before echocardiographic examination. Therefore, SHHF-TMAO (n=8) and SHHF-controls (n=6) were included for further analysis.

**Fig. 11.**
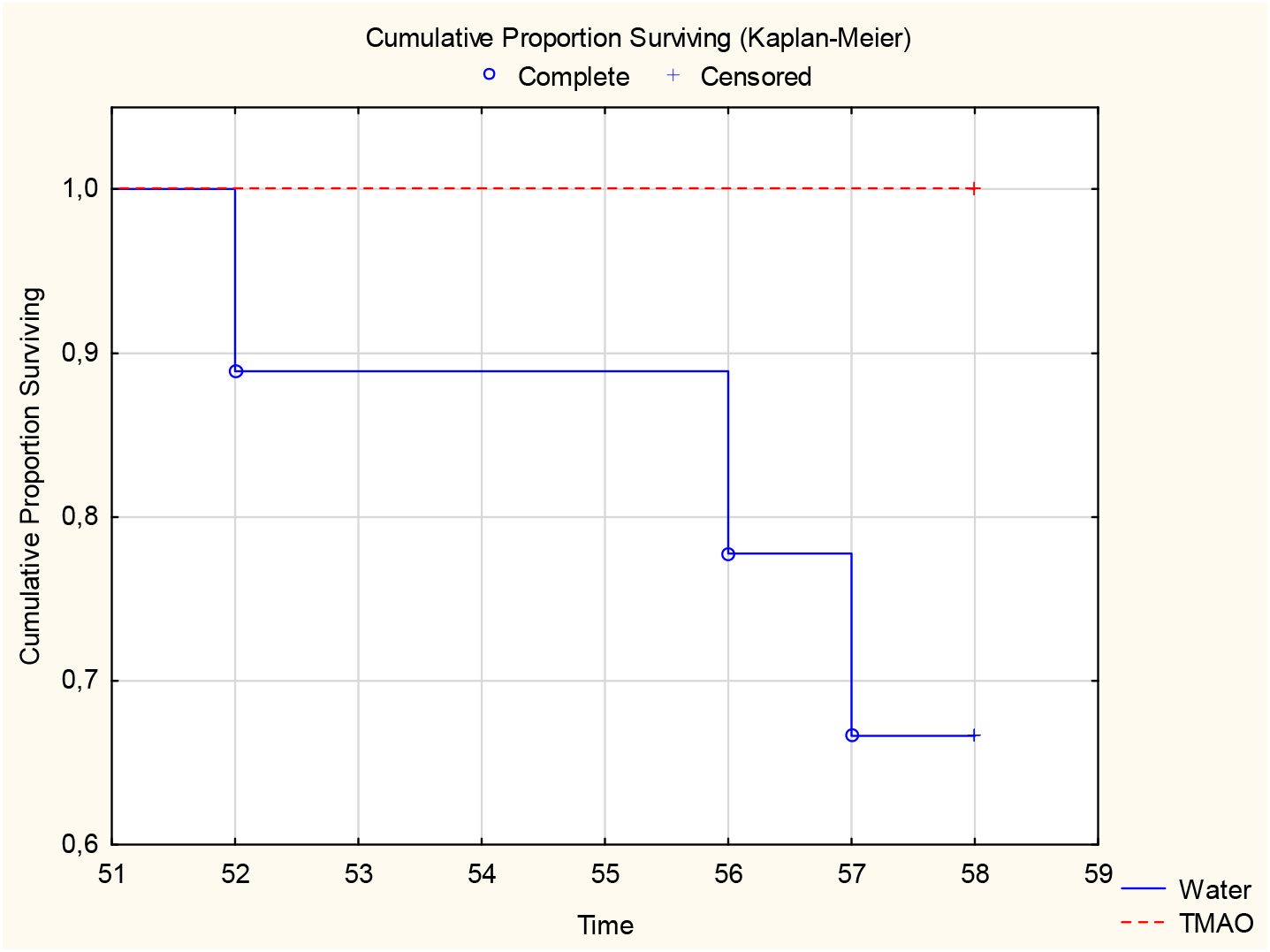
Survival Kaplan-Meier curves for SHHF -- Spontaneously Hypertensive Heart Failure (SHHF/MccGmiCrl-Leprcp/Crl) Rats, treated with either Water (Water) or TMAO in water solution (TMAO). Log-Rank test p = 0.06555

There was no significant difference between groups in food intake. SHHF-TMAO showed 3-4-fold higher plasma TMAO level than Water group. TMAO-treated rats showed significantly higher 24hr urine output and sodium excretion than water-treated rats. TMAO treated group showed significantly lower plasma Ang II level, but there was no difference in aldosterone and vasopressin plasma level (Table 2).

#### Circulatory parameters

Both, SHHF-Water and SHHF-TMAO showed hypertrophic cardiomyopathy with compromised systolic functions and increased plasma NT-proBNP (Fig. 6 and 10, Table 2). However, TMAO-treated rats showed a significantly lower diastolic blood pressure a trend towards a higher stroke volume and ejection fraction (Table 2).

#### Histopathology

The general myocardium pattern and cardiomyocyte morphology in TMAO group did not differ significantly from Water group. Morphometric analysis did not show significant changes in the degree of myocardial fibrosis between the examined groups. However, TMAO-treated rats showed trend towards less myocardial fibrosis (p<0.1), and more visible cardiomyocyte bands and smaller inflammatory infiltration. The histopathological picture of the lungs and kidneys between SHHF-Water and SHHF-TMAO did not differ significantly (Fig. 10).

#### Gene expression

TMAO treated SHHF rats showed significantly lower expression of ATG and AT1R while significantly higher expression of AT2R in the heart. In the kidneys TMAO rats showed a significantly higher expression or renin (Fig. 7 and 8).

### SD-ISO rats: Water vs TMAO treatment

In general, SD subjected to catecholamine stress (isoprenaline) showed some characteristics of Takotsubo-like cardiomyopathy including mild degree of apical akinesis/dyskinesis, edema of cardiomyocytes, increased NT-proBNP level and mild lung edema (Fig. 6 and 12, Table 3, and supplemental Table S1). Numerous, scattered foci of banded mononuclear cell infiltration were visible in the myocardium of SD-ISO rats. Severe hyperemia of myocardial capillaries and arterioles and small organizing foci of myocardial extravasation were present. Nevertheless, the majority of myocardium showed a normal structure. The lungs in the ISO-SD group showed acute pulmonary stasis in the presence of transudate in the alveolar ducts. In the kidneys there was a weak congestion in the medulla and renal bodies. In addition, a small number of tubules filled with an acidic, protein substance. Stimulation of stroma fibrocytes with minimal proliferation without production of connective tissue fibers was observed (Fig. 12).

**Table 3.**
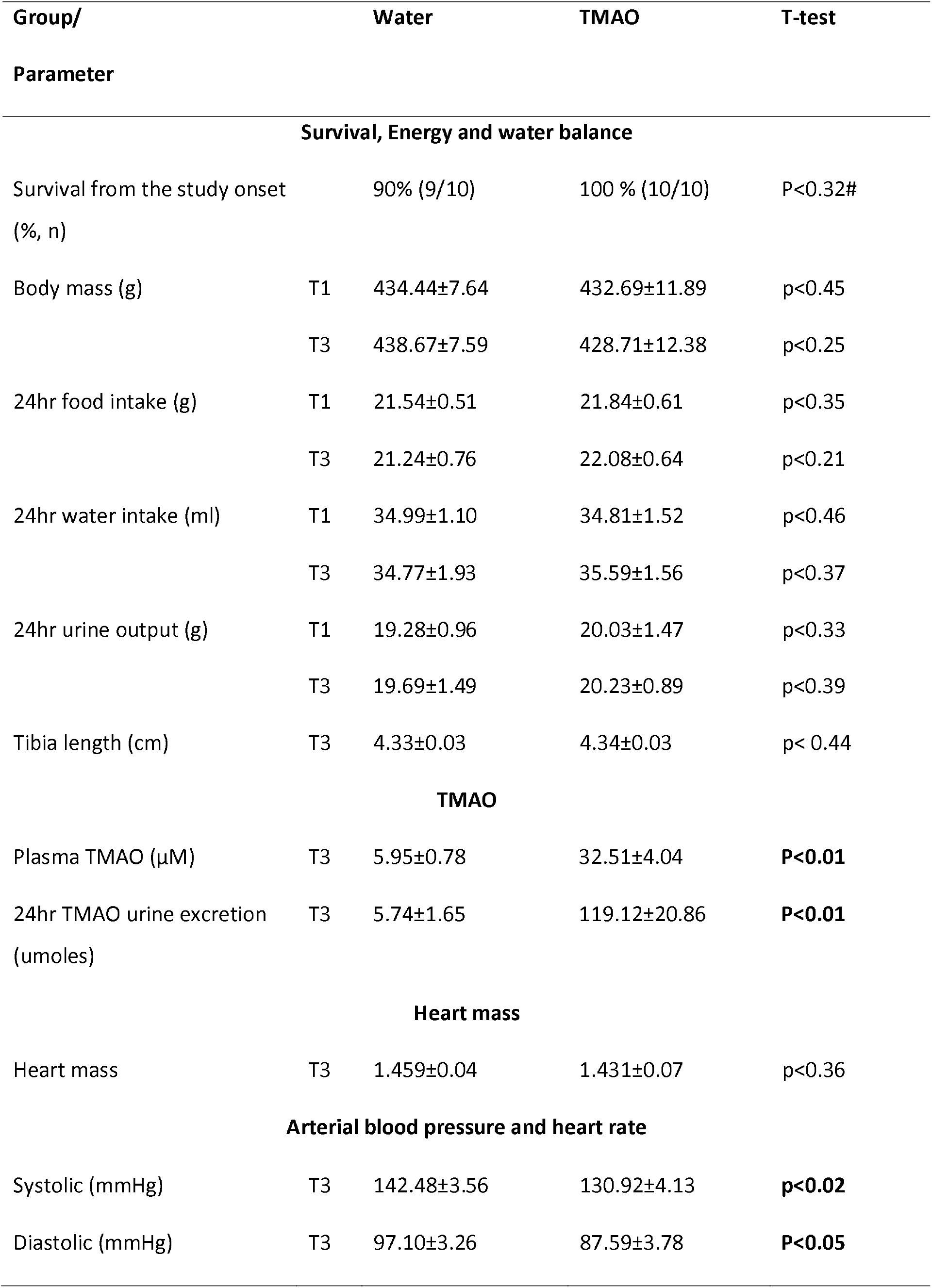
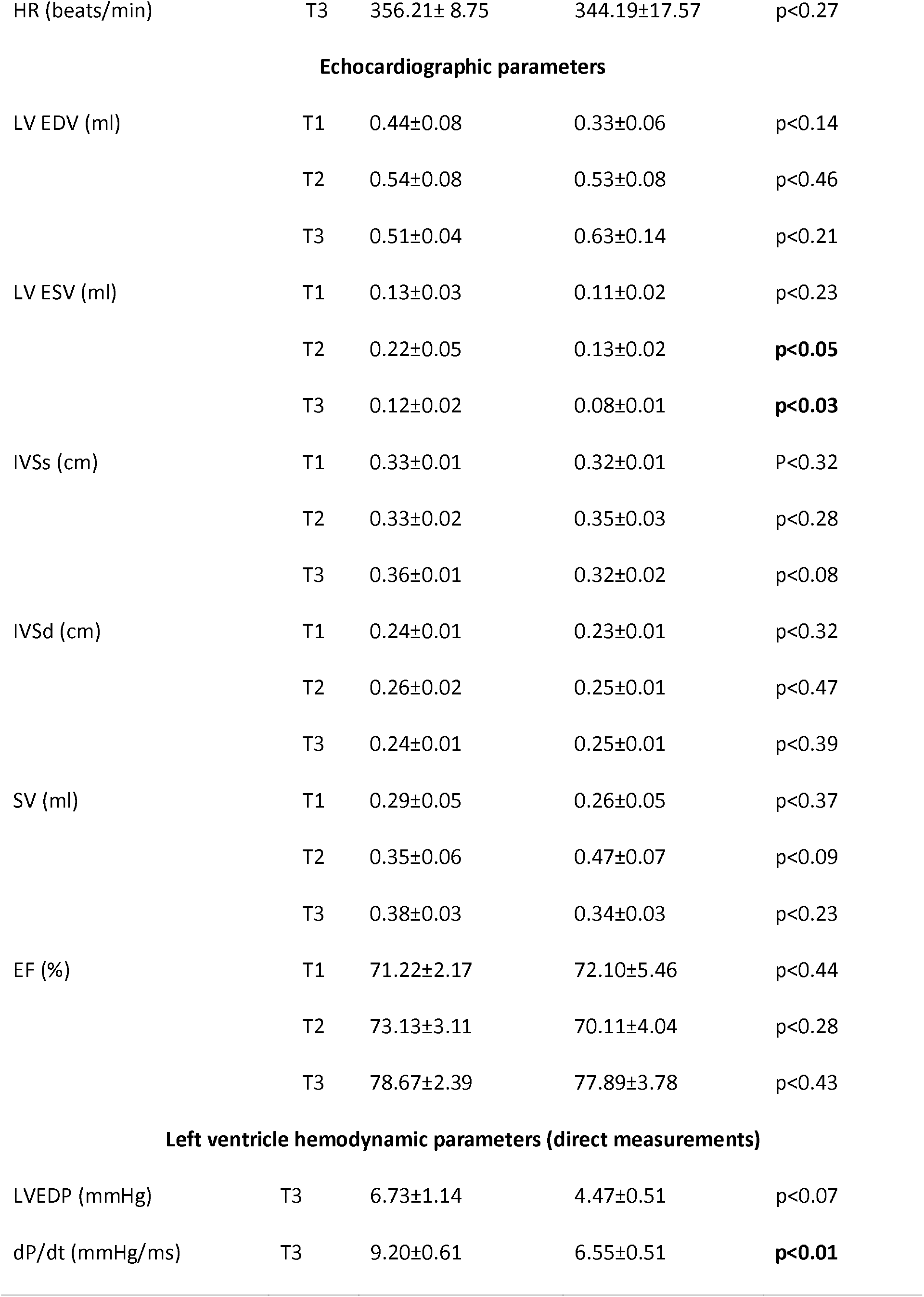
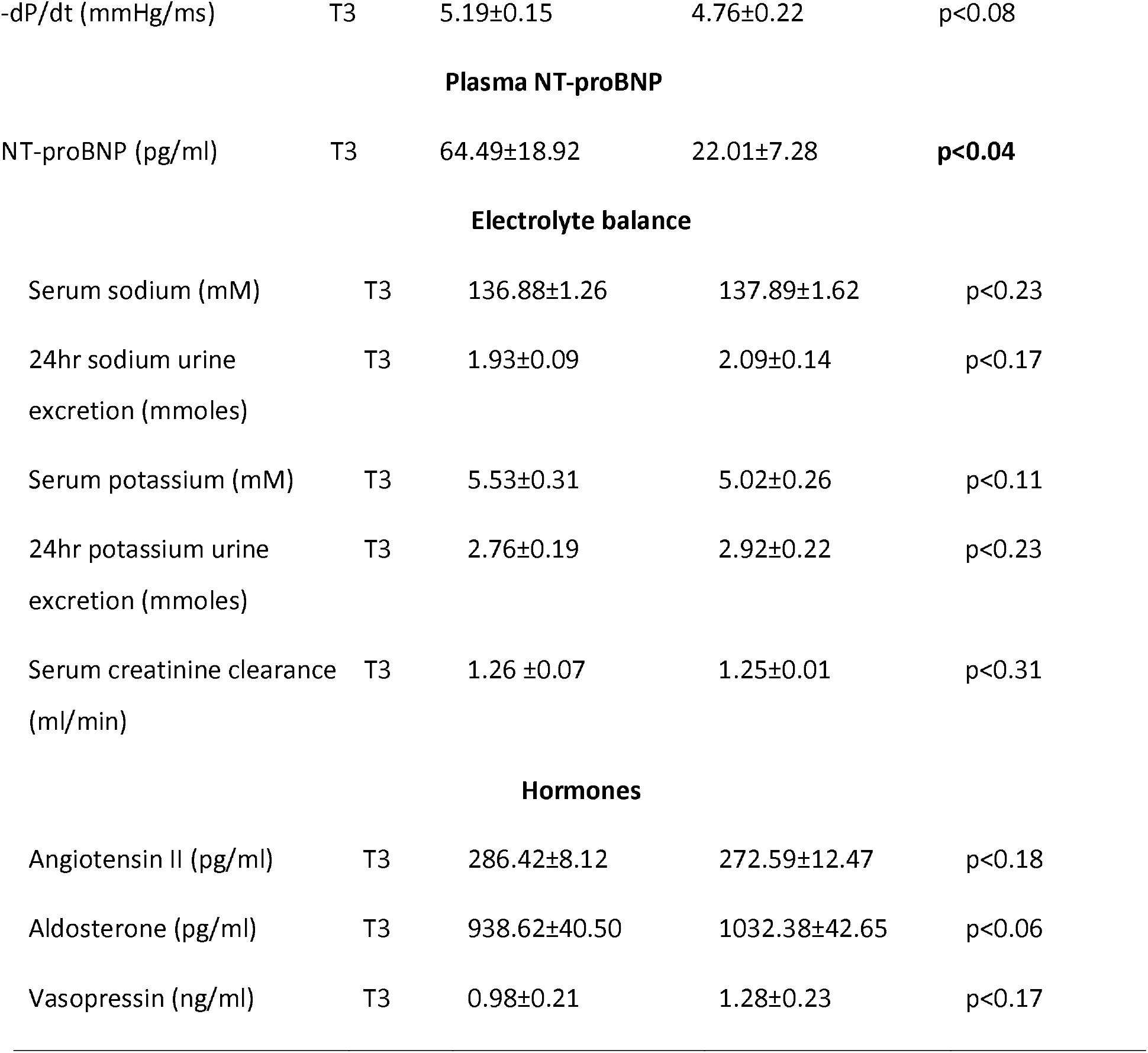
Metabolic, renal and cardiovascular parameters in 58-week-old normotensive Sprague-Dawley rats treated with isoprenaline at the age of 56 weeks. Rats maintained on either Water or TMAO solution in drinking water. T1 - metabolic and echocardiographic measurements, T2 - echocardiographic measurements, T3 - metabolic, echocardiographic and direct hemodynamic measurements (see also the study design, Fig. 1). Creatinine clearance calculated as urine creatinine x urine output (ml/min) / plasma creatinine. LVEDV - left ventricle end diastolic volume, LVESV - left ventricle end systolic volume, SV – stroke volume, EF - ejection fraction, IVSs(d), intraventricular septum diameter during systole and diastole, respectively. LVEDP - pressure in the left ventricle during the end of diastole measured directly with a catheter, +dP/dt - maximal slope of systolic ventricular pressure increment, - dP/dt - maximal slope of diastolic ventricular pressure decrement. Values are means, ± SE. P values by T-test, except # - by log-rank test.

| Group/<br>Parameter |  | Water | TMAO | T-test |
| --- | --- | --- | --- | --- |
| **Survival, Energy and water balance** |  |  |  |  |
| Survival from the study onset<br>(%, n) |  | 90% (9/10) | 100 % (10/10) | P<0.32# |
| Body mass (g) | T1 | 434.44±7.64 | 432.69±11.89 | p<0.45 |
|  | T3 | 438.67±7.59 | 428.71±12.38 | p<0.25 |
| 24hr food intake (g) | T1 | 21.54±0.51 | 21.84±0.61 | p<0.35 |
|  | T3 | 21.24±0.76 | 22.08±0.64 | p<0.21 |
| 24hr water intake (ml) | T1 | 34.99±1.10 | 34.81±1.52 | p<0.46 |
|  | T3 | 34.77±1.93 | 35.59±1.56 | p<0.37 |
| 24hr urine output (g) | T1 | 19.28±0.96 | 20.03±1.47 | p<0.33 |
|  | T3 | 19.69±1.49 | 20.23±0.89 | p<0.39 |
| Tibia length (cm) | T3 | 4.33±0.03 | 4.34±0.03 | p< 0.44 |
| **TMAO** |  |  |  |  |
| Plasma TMAO (μM) | T3 | 5.95±0.78 | 32.51±4.04 | **P<0.01** |
| 24hr TMAO urine excretion<br>(umoles) | T3 | 5.74±1.65 | 119.12±20.86 | **P<0.01** |
| **Heart mass** |  |  |  |  |
| Heart mass | T3 | 1.459±0.04 | 1.431±0.07 | p<0.36 |
| **Arterial blood pressure and heart rate** |  |  |  |  |
| Systolic (mmHg) | T3 | 142.48±3.56 | 130.92±4.13 | **p<0.02** |
| Diastolic (mmHg) | T3 | 97.10±3.26 | 87.59±3.78 | **P<0.05** |
| HR (beats/min) | T3 | 356.21± 8.75 | 344.19±17.57 | p<0.27 |
| Echocardiographic parameters |  |  |  |  |
| LV EDV (ml) | T1 | 0.44±0.08 | 0.33±0.06 | p<0.14 |
|  | T2 | 0.54±0.08 | 0.53±0.08 | p<0.46 |
|  | T3 | 0.51±0.04 | 0.63±0.14 | p<0.21 |
| LV ESV (ml) | T1 | 0.13±0.03 | 0.11±0.02 | p<0.23 |
|  | T2 | 0.22±0.05 | 0.13±0.02 | **p<0.05** |
|  | T3 | 0.12±0.02 | 0.08±0.01 | **p<0.03** |
| IVSs (cm) | T1 | 0.33±0.01 | 0.32±0.01 | P<0.32 |
|  | T2 | 0.33±0.02 | 0.35±0.03 | p<0.28 |
|  | T3 | 0.36±0.01 | 0.32±0.02 | p<0.08 |
| IVSd (cm) | T1 | 0.24±0.01 | 0.23±0.01 | p<0.32 |
|  | T2 | 0.26±0.02 | 0.25±0.01 | p<0.47 |
|  | T3 | 0.24±0.01 | 0.25±0.01 | p<0.39 |
| SV (ml) | T1 | 0.29±0.05 | 0.26±0.05 | p<0.37 |
|  | T2 | 0.35±0.06 | 0.47±0.07 | p<0.09 |
|  | T3 | 0.38±0.03 | 0.34±0.03 | p<0.23 |
| EF (%) | T1 | 71.22±2.17 | 72.10±5.46 | p<0.44 |
|  | T2 | 73.13±3.11 | 70.11±4.04 | p<0.28 |
|  | T3 | 78.67±2.39 | 77.89±3.78 | p<0.43 |
| Left ventricle hemodynamic parameters (direct measurements) |  |  |  |  |
| LVEDP (mmHg) | T3 | 6.73±1.14 | 4.47±0.51 | p<0.07 |
| dP/dt (mmHg/ms) | T3 | 9.20±0.61 | 6.55±0.51 | **p<0.01** |
| -dP/dt (mmHg/ms) | T3 | 5.19±0.15 | 4.76±0.22 | p<0.08 |
| **Plasma NT-proBNP** |  |  |  |  |
| NT-proBNP (pg/ml) | T3 | 64.49±18.92 | 22.01±7.28 | **p<0.04** |
| **Electrolyte balance** |  |  |  |  |
| Serum sodium (mM) | T3 | 136.88±1.26 | 137.89±1.62 | p<0.23 |
| 24hr sodium urine excretion (mmoles) | T3 | 1.93±0.09 | 2.09±0.14 | p<0.17 |
| Serum potassium (mM) | T3 | 5.53±0.31 | 5.02±0.26 | p<0.11 |
| 24hr potassium urine excretion (mmoles) | T3 | 2.76±0.19 | 2.92±0.22 | p<0.23 |
| Serum creatinine clearance (ml/min) | T3 | 1.26 ±0.07 | 1.25±0.01 | p<0.31 |
| **Hormones** |  |  |  |  |
| Angiotensin II (pg/ml) | T3 | 286.42±8.12 | 272.59±12.47 | p<0.18 |
| Aldosterone (pg/ml) | T3 | 938.62±40.50 | 1032.38±42.65 | p<0.06 |
| Vasopressin (ng/ml) | T3 | 0.98±0.21 | 1.28±0.23 | p<0.17 |

**Fig. 12.**
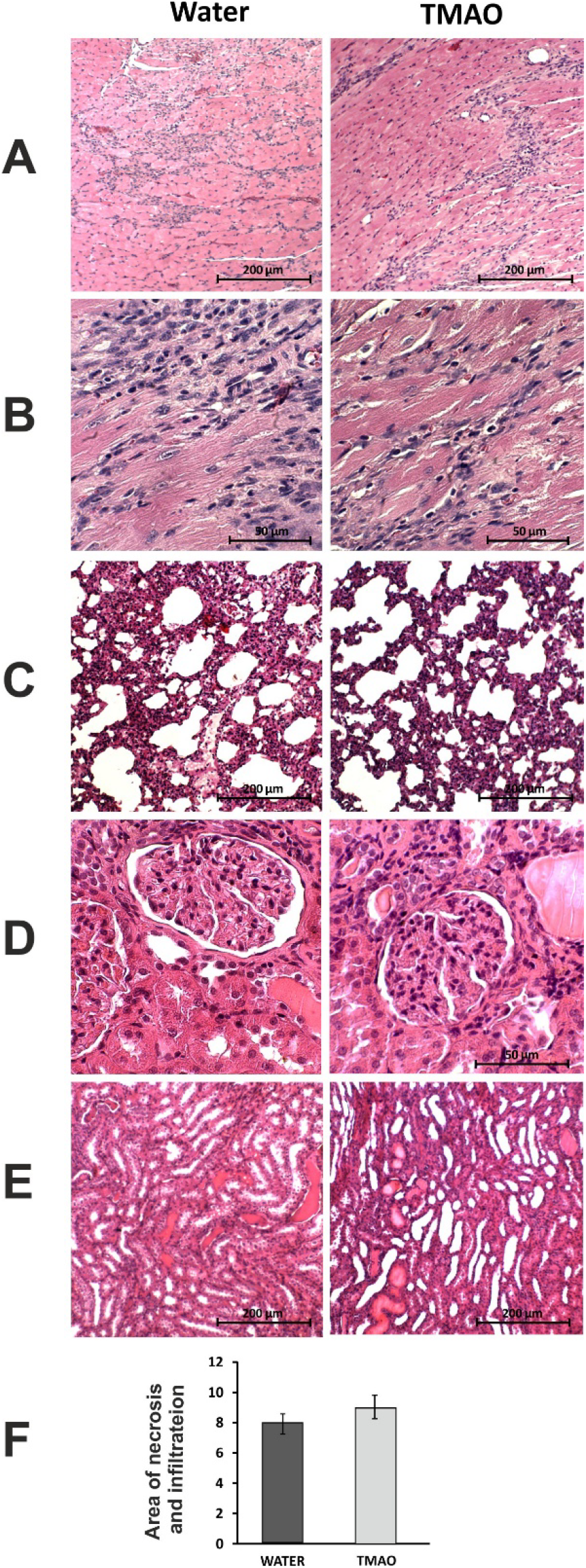
Histopathological picture of heart, lungs and kidneys in Sprague-Dawley rats treated with isoprenaline at the age of 56 week, and drinking either Water or TMAO solution. A - myocardium; haematoxylin-eosin staining at magnification x10; B - myocardium; haematoxylin-eosin staining at magnification x40; C - lungs; haematoxylin-eosin staining at magnification x10; D – kidney - renal cortex, renal bodies; haematoxylin-eosin staining at magnification x40; E – kidney - renal medulla; haematoxylin-eosin staining at magnification x10; F – Percentage of necrotic and inflammatory area in myocardium [%].

#### Survival and water-electrolyte balance

There was no significant difference between ISO-Water and ISO-TMAO groups in survival rate (9/10 vs 10/10, respectively). TMAO-treated rats showed 5 times higher TMAO plasma level than Water-treated rats. There was no significant difference in food and water intake between the groups. There was also no significant difference between the groups in 24hr urine output, however, TMAO treated rats tended to have higher natriuresis (Table 3).

#### Circulatory parameters

TMAO treated rats showed significantly lower systolic and diastolic blood pressure, significantly lower plasma NT-proBNP and lower LV ESV (Table 3).

#### Histopathology

The histopathological picture of the heart, lungs and kidneys between Water and TMAO did not differ significantly (Fig. 12).

#### Gene expression

In the heart and in the kidneys TMAO treated rats showed significantly lower expression of ATG, AT1R. In the kidneys TMAO treated rats showed significantly higher expression of renin (Fig. 7 and 8).

### Effect of TMAO on renal excretion in SD rats - acute studies

The changes in renal excretion induced by TMAO, urea and saline intravenous administration in acute experiments were summarized in Fig. 13. Only the TMAO induced distinct changes in renal excretion. The pattern of diuresis and total solutes excretion induced by TMAO were similar. V and UosmV increases induced by TMAO were associated with transient decrease of Uosm, whereas UNaV and UKV were not affected. This indicates that TMAO did not affected the tubular transport of sodium and potassium but induced osmotic diuresis. The bolus infusions of TMAO and saline produced a transient increase in ABP with no changes in HR, which was followed by a decrease in ABP below the baseline (by 6±3 mmHg). There was no significant correlation between changes in ABP and diuresis.

**Fig. 13.**
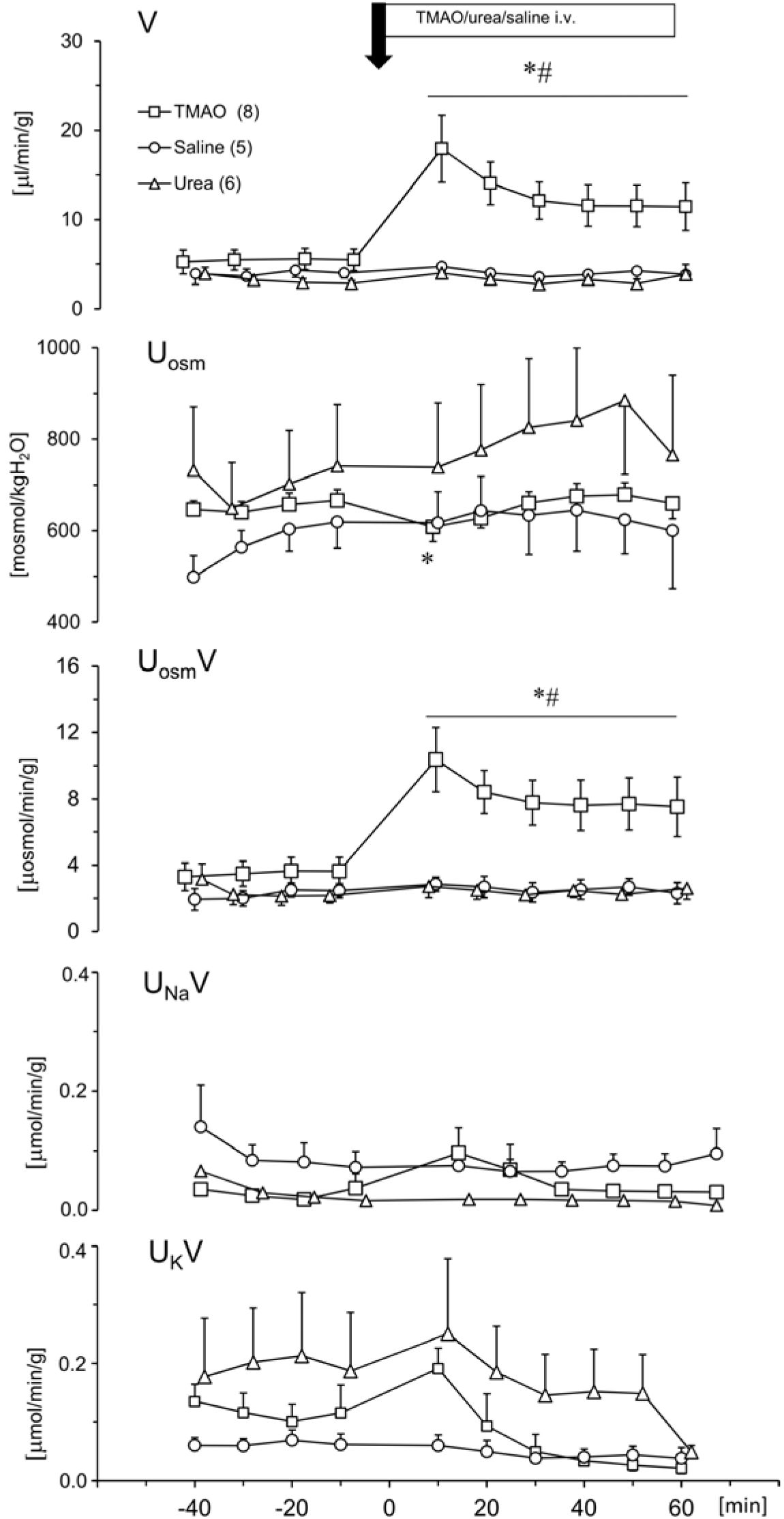
Effects of TMAO, urea and saline on renal excretion in anesthetized Sprague-Dawley rats. The priming dose (indicated by arrow) of TMAO and urea were 2.8 mmol/kg b.w. (bolus), followed by continuous infusion at a rate of 2.8 mmol/kg b.w. / 60 min. V – urine flow; U_osm_ – urine osmolality; U_osm_V, U_Na_V, U_K_V – total solute, sodium and potassium excretion, respectively. Values are means ± SE. * - p < 0.05 vs pretreatment values, ^#^ - p < 0.05 TMAO vs saline, TMAO vs urea.

### Effect of TMAO on structure of LDH exposed to HS and increased temperature

Labeled LDH was stable in PBS solutions, showing no tendency to spontaneous aggregation. The value of diffusion coefficients measured by FCS at 25°C was 49.2 ± 3.3 μm2/s. This corresponds to hydrodynamic radius of around 5.0 nm, which is in line with previously reported values [37].

LDH with and without TMAO subjected to HS (pressure oscillations mimicking those of a rat heart) for 24 hrs did not influence its tertiary or quaternary structure (Fig. 14 A). Similar tests, performed in a different pressure oscillation system, where pressures up to 1000 mmHg were obtained, also brought no observable changes in the protein structure (see the Supplement for details).

**Fig. 14.**
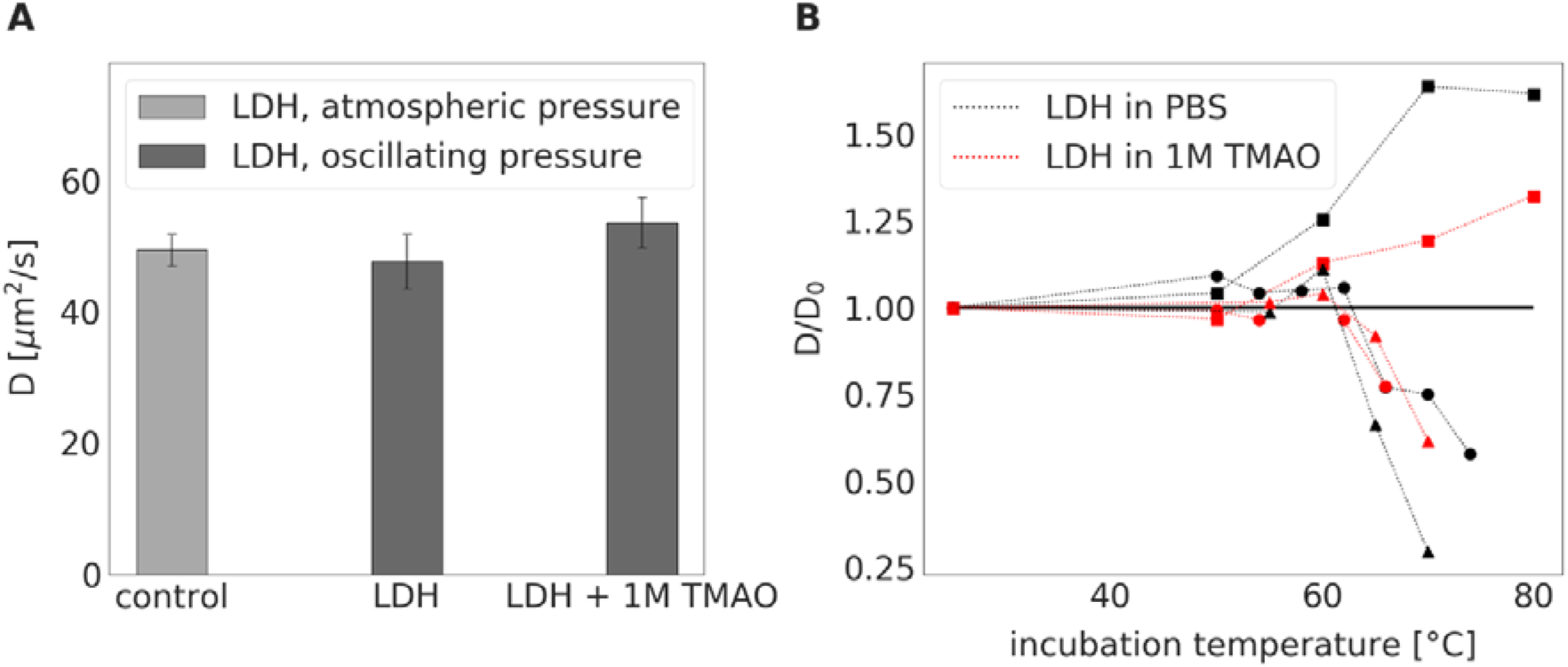
LDH incubated at constant atmospheric pressure for 24 hrs or LDH exposed to pressure oscillation for 24 hrs with or without TMAO. **A** - Diffusion coefficient of LDH either incubated in PBS buffer at constant atmospheric pressure and room temperature (serving as a control) or exposed to pressure oscillation for 24 hrs either in PBS buffer or in 1 M TMAO solution in PBS buffer at room temperature. Irrespective of the presence of 1M TMAO, 24-hour incubation under oscillating pressure did not cause dissociation, denaturation, or aggregation of LDH. **B** – Relative diffusion coefficient (diffusion coefficient, *D* divided by its value in PBS buffer at room temperature, *D*_0_) of LDH exposed to elevated temperatures for 15 min either in PBS buffer (black symbols) or 1 M TMAO solution in PBS (red symbols). Symbol shapes differentiate between three independent measurement series (series 1 – squares, series 2 – circles, series 3 – triangles). Details on this experiment and the complete description of obtained data are presented in the Supporting Information.

The incubation of LDH at atmospheric pressure and elevated temperatures (50 – 80°C) changed the diffusion coefficient of LDH indicating the dissociation of LDH tetramers, as well as its denaturation and aggregation. The addition of 1M TMAO produced a moderate stabilizing effect on LDH, i.e. shifted the threshold of observed protein morphology change towards higher temperatures and mitigated the magnitude of these changes (Fig. 14 B, see also the supplement).

## DISCUSSION

Our study provides evidence that TMAO exerts beneficial effect in HF rats. The beneficial effect of TMAO is associated with diuretic, natriuretic and hypotensive actions rather than a stabilizing effect on cardiac LDH. Specifically, we found that HS at the magnitude substantially greater than that generated by the contracting heart was neutral for cardiac LDH protein structure, with and without TMAO.

Plasma TMAO originates from trimethylamine (TMA), a gut bacteria product, which is oxidized by the liver to TMAO, and from dietary TMAO-rich seafood [29, 36]. Several years ago Tang and collaborators showed that increased plasma TMAO is associated with an increased risk of major adverse cardiovascular events [25], which generated numerous clinical and experimental research suggesting negative effects of TMAO (for review, [15, 36]), However, not all clinical studies have confirmed association between high plasma TMAO and increased cardiovascular risk [14, 24, 34].

Our previous studies suggests that TMAO is only a surrogate marker and that the toxic cardiovascular effects are caused not by TMAO, but by TMAO precursor i.e. TMA [8, 9, 11]. Notably, the latter is a well-established toxin [18]. Since TMA is oxidized (likely, detoxified) by the liver to TMAO, therefore the increased plasma TMAO is simply a proxy of the plasma level of toxic TMA. What is more, recent experimental studies provide evidence for beneficial action of TMAO at low doses [2, 4, 7].

Here, we investigated the effect of TMAO in (i) healthy SD rats, (ii) SHHF (SHHF/MccGmiCrl-Leprcp/Crl) which showed characteristics of heart failure with compromised systolic function and (iii) SD subjected to catecholamine stress (isoprenaline). Our findings show that increased dietary TMAO which produces 3-5-fold increase in plasma TMAO exerts favorable effects on the circulatory system in HF rats. In this study, during a stress-free 52-week observation fatality rate in SHHF treated with TMAO was 0 in comparison to 33% in the untreated group. This was associated with several potentially beneficial hemodynamic and biochemical characteristics in TMAO treated rats such as lower diastolic BP, higher diuresis and natriuresis, lower expression of ATG and AT1R in the heart with concomitant increase in AT2 receptors, and a trend towards lower cardiac fibrosis. Regarding the latter the tissue angiotensin system significantly contributes to heart remodeling and its inhibition is critical in the treatment of heart failure [12]. It is worth stressing that due to high mortality in the Water group the final analysis of hemodynamic and biochemical parameters included only the survivors i.e. the healthiest rats. If all water-treated rats have survived till the end of the experiments the differences between TMAO and Water groups would likely be greater.

The above chronic experiments imply that the beneficial effect of TMAO was dependent on increased diuresis and natriuresis, which is a cornerstones of HF treatment. We assumed that the diuretic and natriuretic effects of TMAO resulted from osmotic diuresis. To evaluate the notion we did additional acute experiments in rats. We found that intravenous administration of TMAO, but not saline given at the same volume, significantly increased diuresis in anesthetized rats. The findings of the acute experiments together with the results of the chronic studies provide strong evidence for a significant diuretic effect of TMAO. Besides, in chronic studies TMAO-treated rats showed increased expression of renin in the kidneys which is a characteristic response to osmotic diuresis which is associated with decreased concentration of sodium in filtrate reaching distal convoluted tubule.

Interestingly, there is evidence that the prevalence and mortality rates of HF are relatively lower in Japan in comparison with the USA and Europe, despite the fact that Japan has higher proportion of older people in the world [17]. At the same time, Japanese have been found to have a significantly higher urine TMAO concentrations than Americans [6], which seems to result from high dietary intake of TMAO-reach seafood by Japanese population.

TMAO, similar to urea, is very osmotically active agent that plays a role of an osmolyte in animals [32]. Notably, before the development of thiazide diuretics, the diuretic treatment of advanced HF comprised of urea infusions [5, 23].

Importantly, in our experiments at the same molar doses TMAO produced significantly higher diuretic response than urea.

A number of biological and biophysical studies indicate that TMAO is not only osmolyte but also a piezolyte, i.e. stabilizes structural proteins and enzymes such as LDH in conditions of increased hydrostatic pressure [20, 33, 35]. This property of TMAO is used by marine animals exposed to high hydrostatic pressure (deep water). Therefore, deep-water animals accumulate TMAO to protect cell proteins from HS [32, 33].

In a healthy heart under physiological conditions the hydrostatic pressure may change between diastole and systole from 0 to 120-130 mmHg. In hypertensive heart or in the heart exposed to catecholamine stress, a diastole-systole change may exceed 220 mmHg, with a frequency of up to 200/min in humans, and higher in small animals. This produces environment in which hydrostatic pressure changes 100 000 times per 24 hours.

Based on several studies showing that TMAO stabilizes teleost and mammalian lactate dehydrogenases against inactivation by hydrostatic pressure [1, 31, 33] we hypothesized that TMAO may exert protective effect on the circulatory system by stabilizing cardiac LDH, a complex enzyme that plays essential role in cardiac metabolism [28].

To evaluate the above hypothesis we have built a unique experimental setup to mimic changes in hydrostatic pressure that occur in the heart under conditions of catecholamine stress. The experiments showed that changes in hydrostatic pressure generated by the contracting heart, and even much greater, are neutral for protein structure of cardiac LDH. Finally, the addition of TMAO did not produce any effect, suggesting neither positive nor negative effect of TMAO on LDH exposed to HS. Nevertheless, we showed that TMAO stabilized LDH exposed to other denaturant, i.e. high temperature, which is in line with other studies [22] and confirms stabilizing action of TMAO on proteins.

Altogether our study for the first time shows that ingestion of TMAO exerts beneficial effect in cardiovascular pathologies associated with fluid retention such as HF. The beneficial effect of TMAO seems to results from its diuretic action. In contrast, the protein stabilizing effect of TMAO is rather not involved in the protection of cardiac proteins against HS produced by the contracting heart. First, we found no effect of TMAO and HS similar to that generated by the heart on protein structure of cardiac LDH. Second, the beneficial effect of TMAO was most pronounced in SHHF whereas it was less evident in SD exposed to catecholamine stress and virtually not present in healthy rats, however, also the latter two groups showed higher diuresis or natriuresis and lower expression of ATG and AT1R in the heart.

### Limitations

A limitation of our study is that biochemical and hemodynamic measurements were performed only at the end of the treatment. This is because we aimed to avoid stress-related circulatory complications in SHHF which are very prone to lethal cardiovascular events.

### Conclusions

TMAO, a molecule present in seafood and a derivate of gut bacteria metabolism, exerts beneficial effect in HF rats which is associated with diuretic, natriuretic and hypotensive effects. The magnitude of hydrostatic stress generated by the contracting heart is neutral for cardiac LDH protein structure. Further studies designed to evaluate TMAO-dependent diuretic and natriuretic effects are needed.

## ELECTRONIC SUPPLEMENTARY MATERIAL

Supplemental Methods and Results

Supplemental Tables

Supplemental Figures S1-S7

## DECLARATIONS

### Ethics approval

The study was approved by the Local Bioethical Committee in Warsaw (permission:100/2016 and 098/2019).

### Consent to participate

Not applicable.

### Consent for publication

Not applicable.

### Availability of data and material

Data are available from the corresponding author upon request.

### Code availability

Not applicable.

## Acknowledgments

The authors are grateful to Tomasz Hutsch MDV, PhD, a veterinary pathologist, for consultations on histopathological analysis.

## Authors’ contributions

Study conception Marcin Ufnal. Material preparation, data collection and analysis were performed by Marta Gawrys-Kopczynska, Marek Konop, Klaudia Maksymiuk and Marcin Ufnal for chronic animal studies, Marta Gawrys-Kopczynska for molecular biological studies, Katarzyna Kraszewska, Ladislav Derzsi, Krzysztof Sozański, Robert Holyst and Marta Pilz for studies with oscillatory-pressure controller and fluorescence correlation spectroscopy, Emilia Samborowska for biochemical analysis, Leszek Dobrowolski, Kinga Jaworska, Izabella Mogilnicka and Marcin Ufnal for acute animal studies. The first draft of the manuscript was written by Marcin Ufnal and all authors commented on previous versions of the manuscript. All authors read and approved the final manuscript.

