## supplement for "TMAO, a seafood-derived molecule, produces diuresis and reduces mortality in heart failure rats"

**Basic Research in Cardiology**

Marcin Ufnal, MD, PhD

Department of Experimental Physiology and Pathophysiology, Medical University of Warsaw

Banacha 1B, 02-097 Warsaw, Poland

­

**SUPPLEMENTAL METHODS AND RESULTS**

**Description of the oscillatory-pressure controller design for the study and correlation spectroscopy of proteins.**

1. **Systems for generation of pressure oscillations**

The setup consisted of two main parts: i) a custom-built oscillatory pressure controller with solenoid micro valves to control the inner pressure and “pulse” frequency and ii) sample chamber. We designed and constructed 3 different samples chambers, which allowed us to expose the samples to pressure oscillation in different ways. We put protein solutions with and without 1M TMAO into the sample chamber and set the pressure values and oscillation frequencies similar or higher than the heart rate and systolic/diastolic pressures of the SHHF rats (i.e. about 200-400/min and 0/200 mmHg, respectively).

a. **Oscillatory pressure controller**

The custom-built oscillatory-pressure controller consisted of two pressure sources with constant but different air pressures (*p1* and *p2*, respectively), each connected through a plunger-type solenoid microvalve (V165, Sirai, Italy) to the inlet/outlet of the sample chamber via teflon tubes (ID/OD=0.8/1.6 mm, Bola, Germany) as shown in Fig. S1. The two microvalves were controlled by a multiplexer switch module (National Instruments, USA) interfaced by a custom-made LabView software. Due to their electro-magneto-mechanical construction the microvalves had some responsive time: the delay between application of the current and opening of the valve was 24 ms, and 8 ms between stopping the current and shutting the valve [1]. The time shifts were taken into consideration while calculating the required oscillatory pressure.

**Operation protocol:** The two valves were initially closed and the pressure inside the sample chamber was equal to the atmospheric pressure. Initiating the oscillatory system, first the first valve opened for 50 ms. As the first valve closed, the second one opened for 50 ms followed by a 50 ms pause when both valves were closed. With the response times included, one full cycle took 214 ms, generating an oscillation of ~280/minute between pressure *p1* and *p2*, mimicking the heart-beat of a SHHF rat. In each of our experiments we used this kind of pressure-oscillatory system, only the applied pressures *p1* and *p2* were different.

b. **Microfluidic heart chip preparation.**

In the first set of experiments a microfluidic device was constructed in polydimethylsiloxane (PMDS; Sylgard 184, Dow Corning, USA) in three steps: first the channels and sample chamber (in the shape of a heart) were micro-milled in a polycarbonate plate (PC; Macrolon, Germay) using a CNC milling machine (MSG4025, Ergwind, Poland). Then, we poured PDMS onto the PC chip, polymerized the PDMS at 75°C for 2 hours, activated the surface by Laboratory Corona Treater (BD 20AC, Electro-Technic Products, USA) and silanized in the vapours of tridecafluoro-1,1,2,2-tetrahydrooctyl-1-trichlorosilane (United Chemical Technologies, USA) for 60 min under vacuum (10mbar). Finally, this negative mold was used to produce the final PDMS chip. Inlet and outlet holes were punched using a small diameter (ID=0.8mm) biopsy puncher prior to bonding the PDMS chip to 1mm thick glass slides using oxygen plasma for 45s.

**Operation parameters.** The inlets were connected to the heart-shaped sample chamber (height of 300 µm and total volume of ~220µL) via microchannels from two sides (Fig. S2a.). On the one side the microchannel ended in a single inlet through which the sample liquid was injected. Once the chamber was completely filled the inlet was sealed air-tight. On the second side of the chamber the microchannel branched into two, ending in 1-1 inlets which were connected to the two pressure sources (*p1* and *p2*) of the oscillatory system by Teflon tubes (ID=0.8mm, OD=1.6mm; Bola, Germany). The valves were initially closed and inside the chamber atmospheric pressure acted on a sample liquid.

In the first run we applied *p1*=250 mmHg overpressure and *p2*= 0 mmHg (i.e. atmospheric pressure) and oscillation frequency of ~280/minute as described above.

**Results.** The inlet channel (from the side of the oscillatory system) was filled with the sample liquid only half way. As the pressure was oscillating in the microfluidic device, rapid movement of the liquid meniscus could be observed according to the oscillation frequency (see Fig 2b-d). The overpressure *p1* pushed the liquid meniscus towards the chamber, while upon switch to the low pressure (p2) the meniscus pulled back. Unfortunately, since the liquid was in direct contact with the oscillating air some evaporation occurred. As a consequence, the sample liquid slowly evaporated from the chamber and in ca. 40 minutes from starting the experiment and the microfluidic chamber dried out completely. Various pressures and oscillation frequencies were used but in each case the sample evaporated within 30-60 minutes.

c. **PDMS sample chamber in the shape of a micro centrifuge-tube preparation.**

To avoid evaporation we constructed a different type of microfluidic device, where the sample liquids were not in direct contact with the oscillating air. We constructed a different type of microfluidic device as follow: we removed the caps of four 0.5 mL conical tip micro centrifuge tubes (Eppendorf, Germany) and glued them to a microscope glass slide (75x24x1 mm) in line, separated by ~3 mm from each other. On two other, larger glass plates (75x50x1 mm) we glued small blocks of polycarbonate (45x15x5 mm) and fixed them from two sides of the micro centrifuge tube array, so that the distance between the polycarbonate blocks and the tubes was from both sides ~1mm. We put this construction into a small box, filled it with polydimethylsiloxane (PDMS elastomer, Sylgard 184 mixed with curing agent in ration 10:1 and degassed) and polymerized the PDMS at 75°C for 2 hours. After that, we removed it from the container, removed the glass plates, the polycarbonate blocks and micro centrifuge tubes from the cured PDMS. Next, we bonded the PDMS block from the two sides where the removed polycarbonate blocks left a cuboid-shaped cavity to 1 mm glass plates (75x50x1 mm) using oxygen plasma. We also inserted steel capillaries (OD = 0.8 mm) from two sides to each cavity, to which we connected the oscillatory system by Teflon tubes (Bola, Germany). Next, we constructed a slab of PDMS (thickness of 4 mm) and used it as the cover plate on the top of the chamber array which was left after the Eppendorf tubes were removed from the PDMS. Prior to bonding with oxygen plasma we punched holes for sample injection on the PDMS slab. Once the microdevice was ready (see Fig. S3.d) we injected the sample liquids through the punched inlet holes into the sample chambers and sealed the inlet holes air tight. The sample liquids filled the chamber completely. In a separate experiment we left small air bubbles in the Eppendorf-chambers for pressure estimation (see below).

**Operation parameters**. After the chambers were filled the oscillatory system was turned on using pressures *p1* = 2.5 bar and *p2* = 500 mbar (equivalent to 1875 and 375 mmHg, respectively). Higher pressure than 2.5 bar resulted in destruction of the PDMS membrane between the sample chambers and pressurized cavity. As the high pressure air filled the cuboid-cavities, they expanded and pushed the 1 mm thick PDMS membrane towards the sample-filled chambers squeezing and deforming them (on the other side of the cavity the glass avoided the expansion). As the valve of the high pressure source (*p1*) closed and the second valve opened the pressure in the cavity significantly dropped and the deformed sample chambers returned to their original shape. The rapid oscillation of the air pressure (~280/min) in the cavities resulted in corresponding squeezing-releasing pulsation of the chambers, mimicking even more a beating heart than the previous method. After 24 hours of operation, the system was turned off, the plugs were removed from the inlet holes and the processed samples for analysis were removed with a syringe and needle.

**Comments.** The disadvantage of this method was that it was not possible to directly measure the exact pressure increase inside the sample chambers. We have estimated it indirectly by measuring the change of the diameter of some small, trapped bubbles in the sample liquid. From this, we calculated the volume change of the bubble and used Boyle’s law (*pV=const.*) to calculate the corresponding pressure change. We have estimated the pressure increase as approximately 180-250 mmHg.

d. **Pressure-bottle-pressure significantly exceeding the physiological range preparation.**

We put 4 samples into four 0.5 mL micro centrifugal tubes (Eppendorf tubes), removed the caps of the tubes and covered them with a parafilm (Bemis, USA) instead. Small holes were made in the parafilm with a syringe needle (OD = 0.6 mm). We put the sample-loaded Eppendorf tubes inside a 250 mL laboratory glass bottle (Duran™, Fischer Scientific, Germany) and closed air tight with a screw cup. Before closing two small holes in the screw cap were drilled, steel capillaries (OD = 0.8 mm) were placed in and glued with epoxy-based adhesive. The oscillatory system was connected to the steel capillaries with Teflon tubes.

**Operation parameters**. In this experiment we decided to work well above the target pressure range, in order to increase the chance for protein denaturation. After few pilot-experiments we set the high pressure *p1* equal to 4.5 bar (3375 mmHg) and the lower pressure *p2* equal to 0.5 bar (375 mmHg). However, due to the relatively large volume of the bottle, high compressibility of the air and the short opening times of the two valves the pressures acting the samples were not equal to the input pressures *p1* and *p2*.

To measure the actual pressure range inside the laboratory bottle, which acted directly on the samples, we attached two manometers (model 82100, AZ Instruments, Taiwan) to out setup: one before the second valve to measure the pressure that builds up inside the laboratory bottle and another behind the second valve to measure the pressure after release.

The manometers on the other hand had much longer response times (500 ms) than the valve’s opening-closing times. As a result, the manometer indicated constantly varying values but over a longer period of monitoring we noticed that these values always oscillated between a specific range. Therefore, despite all the technical difficulties, with careful monitoring over the period of some 15-20 minutes we were able to conclude the following:

i) The pressure in the laboratory bottle increased continuously for the first 50-60 seconds before reaching its maximum. This indicates that the first valve allowed more air to flow in than the second valve let flow out (even the opening times for both valves were set to 50 ms). This is probably due to the fact that the response time of the second valve was somewhat faster than of the first valve.
ii) We also observed some periodic desynchronization and re-synchronisation of the two valves: each 15-20 second the first manometer indicated values between 1600 and 1000 mbar, then dropped to values between 1200 and 750 mbar, then increased again. This also indicates the faster response time of the second valve.
iii) During the ~15 minutes monitoring, the first manometer never showed lower pressure than 700 mbar. The maximum value observed was around 1600 mbar.
iv) The lowest pressure the second manometer indicated was 450 mbar.

We can conclude that in the laboratory bottle the pressure acting on our samples never dropped below 450 mbar (320 mmHg) and increased up to 1600 mbar (1200 mmHg). The oscillation of the pressure was ~230/min with some “desynchronization” each 15-20 s when the pressure difference *Δp=p1-p2* was only ~150 mbar (110 mmHg) and with pressure difference of ~600 mbar (450 mmHg) during synchronized operation. We notice, that the manometer also had a dead volume of few milliliters, thus the real pressure and the range in between it oscillated could be even higher.

We left the samples in the pressurized laboratory bottle under these conditions for 24 hours, then depressurized the bottle and took out the samples for analysis.

2. **Fluorescence correlation spectroscopy: experimental details**

Fluorescence correlation spectroscopy (FCS) measurements were performed using a system based on a Nikon C1 inverted confocal microscope. Water immersion Nikon PlanApo 60x objective with a correction collar was used. Excitation was done using LDH-series diode lasers by PicoQuant, driven by a Sepia II controller. Pulsed laser excitation and fluorescence lifetime filtering was applied to reduce possible artifacts in the autocorrelation. Laser powers in the range of several microwatts were used to prevent photobleaching and reduce triplet state contribution. A complete time-correlated single-photon counting system by PicoQuant with an avalanche photodiode by Perkin-Elmer was fitted to the microscope for photon detection. Computer-controlled environmental chamber by Oko-lab was used to maintain a temperature of 25°C during all FCS measurements.

Throughout the experiments, we used Atto 488 and Atto 647N dyes for labeling the proteins (the dyes differ in charge and hydrophobicity, which may somewhat affect the physicochemical properties of the labeled protein – however, we did not find any significant difference in either diffusion coefficients or tendency to aggregation or dissociation of the studied protein when using the two different dyes). For Atto 488 excitation a 485 nm laser was used and calibration was done with rhodamine 110. For Atto 647N excitation a 636 nm laser was used, with calibration done using Alexa 647.

Protein was labeled using active NHS esters of the respective dyes, according to a protocol supplied by the manufacturer. A 10-fold excess of the dye was used to ensure high degree of labeling (which was especially important for LDH, where we intended to have at least one label per subunit to be able to monitor probe concentration changes upon dissociation of the tetramer). Post-labeling purification was performed using Bio-Rad BioGel P-30 size exclusion columns. Protein labeling and purification was performed immediately before starting each experimental run.

For all FCS experiments, directly prior to the measurement, all samples were diluted to a concentration in the nanomolar range, which is optimal for this technique. The solvent was PBS with addition of 0.002% Tween20 surfactant. The surfactant was used to discourage adsorption of the proteins on the sample chamber walls as well as sample/air interface, which would preclude quantitative concentration measurements. The concentration of Tween 20 of 0.002% corresponds to a molar concentration that of ~16 μM, whereas the critical micellization concentration is ca. 60 μM (~0.0074% by weight). Therefore, since there were no supramolecular structures of the surfactant present in the solutions and the surfactant was non-ionic, it should not influence the structure of the proteins suspended in the solution [2].

a. **FCS results: proteins under pressure higher than physiological**

Figure S5 summarizes the FCS measurement results of LDH samples incubated at non-physiologically high pressure (up to 1200 mmHg peak) with pulse-like pressure oscillations. Sample incubation was conducted over a period of 24 hours at 25°C, using pressure oscillation system describe above (Fig. S3). LDH concentration during incubation was around 1 μM; FCS measurements were performed after 100-fold dilution in PBS supplemented with Tween20. The recorded diffusion coefficients are, within the experimental error, indistinguishable between the sample subject to pressure treatment and control. Similarly, presence of TMAO does not contribute any change to the observed result.

b. **FCS results: proteins under thermal stress**

While we did not observe any change in the FCS results for LDH upon exposure to pressure oscillations, we decided to perform an additional in vitro experiment with a different stress factor, i.e. elevated temperature. The procedure for these experiments was as follows:

i) LDH solutions, in either pure PBS or PBS supplemented with 1M TMAO, were pipetted into Eppendorf tubes;
ii) samples were placed in a water heat bath (Lauda, electronically controlled) and incubated for 15 minutes (in separate tests – data not shown here – we verified that prolonging the incubation time above this limit, up to 1 hour, did not influence the obtained results);
iii) then, samples were immersed in room-temperature water bath to cool down;
iv) FCS measurement was performed for samples equilibrated to 25°C.

Figure S6 compiles results of 3 independent repetitions of such experiments; each series involved LDH in pure PBS and PBS supplemented with 1M TMAO. Diffusion coefficients are rescaled by the diffusion coefficient recorded for a given sample prior to being subject to elevated temperature. This is to eliminate the influence of TMAO on the viscosity of the solution, which directly affects the diffusion coefficients, so that the effects related directly to changes in the protein structure can be directly visualized.

In the first experimental series (data marked as squares in Figure S6), an increase of the average diffusion coefficient of LDH by a factor of 1.30 and 1.65 (with and without TMAO, respectively) was noted at temperatures above 70°C. Native LDH is a tetramer. In a first approximation (spherical particles), dissociation of a tetramer to monomers should cause a decrease of the observed *D* by a factor of around 1.6. We can therefore suppose that we indeed observed dissociation of the tetramer to monomers. However, it must be taken into consideration that FCS measurement result is an ensemble average of all the fluorescent species present in the solution, while discriminating between various subpopulations of fluorescent species of similar sizes may not be possible. Thus, it cannot be ruled out that also protein denaturation was happening at the same time and the obtained result contains contributions from various subpopulations present in the sample (tetramers, folded monomers and unfolded monomers). No aggregation was observed in this case.

In the two following experiment series, a slight increase in the diffusion coefficient was observed around 60°C and a sharp drop at higher temperatures. Along with the decrease of *D*, a decline in probe concentration was observed as well as a drop in the autocorrelation quality. This information, combined with raw signal analysis and confocal imaging (data not shown) allows to confidently confirm the formation of large protein aggregate formation. Most probably, this was an aftermath of tertiary structure denaturation.

The sole difference between run 1 (where there was no aggregation) and the following ones (where aggregation was clearly observed) was the initial LDH concentration during sample incubation, which was around 3 nM and tens of nM, respectively. These values are not exact due to possible errors due to adsorption of the protein on the walls of the tubes, tips, etc., which may be substantial at nanomolar concentrations (using an addition of the nonionic surfactant mitigates this effect, but does not fully eliminate it). We hypothesize that in all three experiment series a gradual dissociation of tetramers to monomers was occurring above 55°C, which was followed by denaturation of the tertiary protein structure at higher temperatures. At LDH concentration of the order of tens of nM, aggregation prevailed. At concentration lower by an order of magnitude, aggregation was progressing much more slowly (if at all), while the possibly formed aggregates were too sparse to influence the measurement result. Further work, including probing a broader matrix of LDH concentrations and temperature values, would be needed to confirm this supposition.

It is interesting to note that in all the above experiments the presence of 1M TMAO during LDH incubation shifted the threshold of observed protein morphology change towards higher temperatures and mitigated the magnitude of these changes. Some stabilizing effect of TMAO on the native protein structure may be therefore supposed; however, we do not have enough experimental data yet to propose a detailed mechanism of this effect.

**Histopathological analysis, comparison between SD, SHHF and SD-ISO rats.**

58-week-old, male SHHF - Spontaneously Hypertensive Heart Failure (SHHF/MccGmiCrl-Leprcp/Crl) SHHF, SD – Sprague Dawley rats, SD-ISO: SD rats treated with isoprenaline at a dose of 100 mg/kg at the age of 56 week.

There were no pathological changes in the heart, lungs and kidneys of SD (Fig. S7 – A, D, G, J, M, P).

In SHHF cardiomyocytes, an increase of cell diameter with enlargement of cell nuclei, reduction of cytoplasmic acidity, blurring of striation of cardiomyocytes and intercalated disc were found. The myocardial muscle fiber system was irregular and locally cardimyocytes had a wavy arrangement. Only single cardiomyocytes revealed features of an increased cytoplasmic acidity, nuclei pycnosis, and signs of degeneration. In addition, the amount of stromal connective tissue was increased compared to SD. Weak to moderate mononuclear cell infiltration was seen in the fibrosis and perivascular areas. The described changes were visible in the entire myocardium, but they were particularly severe in the left ventricle. Based on the described pathological changes, we can conclude about dilated cardiomyopathy (Fig. S7 B, E, H)

SHHF lungs showed signs of passive hyperemia, with thickening of the interalveolar septa and a weak focal parenchymal edema of the lungs. Slight stromal connective tissue hyperplasia also were observed (Fig. S7 K).

There were no significant pathological changes in the SHHF kidneys (Fig. S7 N, R)

Numerous, scattered foci of banded mononuclear cell infiltration (formed by lymphocytes and macrophages) are visible in the myocardium of SD-ISO. Stimulated and hyperplastic fibroblasts were also found in these areas but they did not form matrix and fibers of connective tissue. Weak to moderate edema of cardiomyocytes was observed. Moreover, pale pink masses were visible in these foci, which probably arose as a result of liquefactive, fibrillar necrosis of cardiomyocytes, removed by macrophages. Severe hyperemia of myocardial capillaries and arterioles and small organizing foci of myocardial extravasation. Nevertheless, apart from the described foci, the vast majority of myocardium has a normal structure (Fig. S7 C, F, I).

The lungs in the SD-ISO showed acute pulmonary stasis in the presence of transudate in the alveolar ducts (Fig. S7 L).

In SD-ISO weak congestion in the medulla and renal bodies were visible. In addition, a small amount of tubules filled with an acidic, protein epithelial damage and cyst formation. Stimulation of stroma fibrocytes with minimal proliferation without production of connective tissue fibers was observed (Fig. S7 O, S).

**SUPPLEMENTAL TABLES**

**Table S1.** Echocardiographic parameters in 58-week-old normotensive Sprague Dawley rats treated with isoprenaline at the age of 56 weeks. LVEDV - left ventricle end diastolic volume, LVESV - left ventricle end systolic volume, SV – stroke volume, EF - ejection fraction, IVSs(d), intraventricular septum diameter during systole and diastole, respectively. Values are means, ± SE. P values by T-test.

**T1- before the treatment with isoprenaline, T2 -24hrs after the treatment, T3 - 8 days after the treatment.**

|  | SD-ISO-WATER |  | p value |
| --- | --- | --- | --- |
| HR (beats/min) | 323.14±26.56  441.67±8.09  451.10±18.85 | T1 vs. T2  T2 vs. T3  T1 vs. T3 | P <0.0000492  NS ( p = 0.322)  p < 0.0001 |
| LV EDV (ml) | 0.44±0.08  0.54±0.08  0.51±0.04 | T1 vs. T2  T2 vs. T3  T1 vs. T3 | NS (p=0.188)  NS (p=0.340)  NS (p=0.239) |
| LVESV (ml) | 0.13±0.03  0.22±0.05  0.12±0.02 | T1 vs. T2  T2 vs. T3  T1 vs. T3 | NS (p=0.071)  p<0.038  NS (p=0.361) |
| IVVs (cm) | 0.33±0.01  0.33±0.02  0.36±0.01 | T1 vs. T2  T2 vs. T3  T1 vs. T3 | NS (p=0.153)  NS (p=0.086)  NS (p=0.078) |
| IVSd (cm) | 0.24±0.01  0.26±0.02  0.24±0.01 | T1 vs. T2  T2 vs. T3  T1 vs. T3 | NS (p=0.194)  NS (p=0.298)  NS (p=0.353) |
| LVPWs (cm) | 0.38±0.02  0.38±0.03  0.39±0.01 | T1 vs. T2  T2 vs. T3  T1 vs. T3 | NS (p=0.500)  NS (p=0.392)  NS (p=0.356) |
| LVPWd (cm) | 0.25±0.01  0.26±0.01  0.25±0.01 | T1 vs. T2  T2 vs. T3  T1 vs. T3 | NS (p=0.285)  NS (p=0.208)  NS (p=0.397) |
| SV (ml) | 0.29±0.05  0.35±0.06  0.38±0.03 | T1 vs. T2  T2 vs. T3  T1 vs. T3 | NS (p=0.231)  NS (p=0.352)  NS (p=0.095) |
| EF (%) | 71.22±2.17  73.13±3.11  78.67±2.39 | T1 vs. T2  T2 vs. T3  T1 vs. T3 | NS (p=0.319)  NS (p=0.099)  P<0.017 |

|  | SD-ISO-TMAO |  | p value |
| --- | --- | --- | --- |
| HR (beats/min) | 297.25±10.06  440.56±19.93  438.30±15.35 | T1 vs. T2  T2 vs. T3  T1 vs. T3 | p<0.00002  NS (p=0.465)  P<0.000001 |
| LV EDV (ml) | 0.33±0.06  0.53±0.08  0.63±0.14 | T1 vs. T2  T2 vs. T3  T1 vs. T3 | P<0.032  NS (p=0.290)  P<0.038 |
| LVESV (ml) | 0.11±0.02  0.13±0.02  0.08±0.01 | T1 vs. T2  T2 vs. T3  T1 vs. T3 | NS (p=0.458)  P<0.024  NS (p=0.073) |
| IVSs (cm) | 0.32±0.01  0.35±0.03  0.32±0.02 | T1 vs. T2  T2 vs. T3  T1 vs. T3 | P<0.27  NS (p=0.272)  P<0.025 |
| IVSd (cm) | 0.23±0.01  0.25±0.01  0.25±0.01 | T1 vs. T2  T2 vs. T3  T1 vs. T3 | NS (p=0.066)  NS (p=0.0367)  NS (p=0.099) |
| LVPWs (cm) | 0.33±0.01  0.37±0.02  0.36±0.02 | T1 vs. T2  T2 vs. T3  T1 vs. T3 | P<0.049  NS (p=0.394)  NS (p=0.089) |
| LVPWd (cm) | 0.25±0.01  0.26±0.01  0.26±0/01 | T1 vs. T2  T2 vs. T3  T1 vs. T3 | NS (p=0.191)  NS (p=0.361)  NS (p=0.128) |
| SV (ml) | 0.26±0.05  0.47±0.07  0.34±0.03 | T1 vs. T2  T2 vs. T3  T1 vs. T3 | P<0.011  P<0.051  NS (p=0.73) |
| EF (%) | 72.10±5.46  70.11±4.04  77.89±3.78 | T1 vs. T2  T2 vs. T3  T1 vs. T3 | NS (p=0.387)  NS (p=0.094)  NS (p=0.202) |

**SUPPLEMENTAL FIGURES AND FIGURE LEGENDS**

Figure S1. Schematic illustration of the oscillatory pressure controller

**Figure S2.** a) Scheme of the experimental setup with the heart-shaped PDMS microfluidic device. b) time-sequence snapshots of the device in operations.

**Figure S3**. a) Scheme of the set-up with the pressure oscillator connected to the Eppendorf chip with side view and b) top view of the “Eppendorf chip”.

**Figure S4**. Schematic representation of the “pressure bottle” system.

**Figure S5.** Diffusion coefficients,
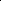
 of LDH measured by FCS.

**Figure S6.** Diffusion coefficients of LDH incubated for 15 minutes at elevated temperatures, scaled by the diffusion coefficient of native (not temperature-treated) LDH.

**Figure S7.** Comparison of the histopathological picture between SD, SHHF and ISO-SD.

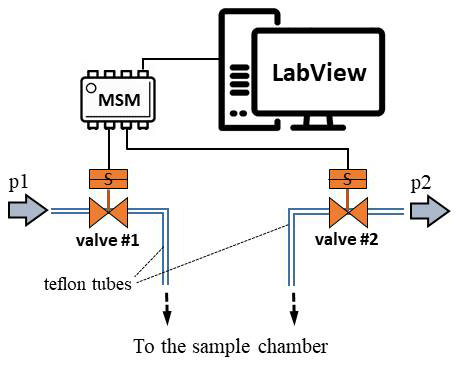

Figure S1. Schematic illustration of the oscillatory pressure controller

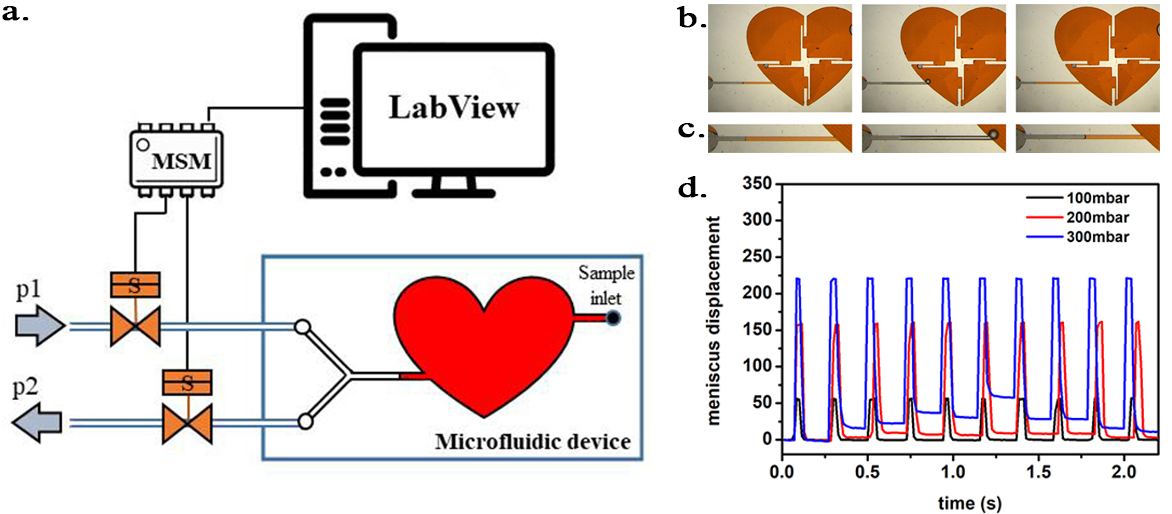

Figure S2. a) Scheme of the experimental setup with the heart-shaped PDMS microfluidic device. b) time-sequence snapshots of the device in operations. From left to right: the device is filled with liquid and is at rest (for better visualisation we used here red-dyed water instead of the transparent protein solution). The microchannel connecting the pressure system to the microfluidic chamber is half-way filled. Applying high pressure (valve 1 open) pushes the liquid meniscus toward the chamber. Closing valve #1 and opening valve #2 (low pressure) the liquid meniscus pulls back (even further than its original position). c) Close-up of the moving liquid meniscus, d) oscillation profile constructed from the position of the liquid meniscus in the microchannel as a function of time for various pressure differences *Δp=p1-p2.*

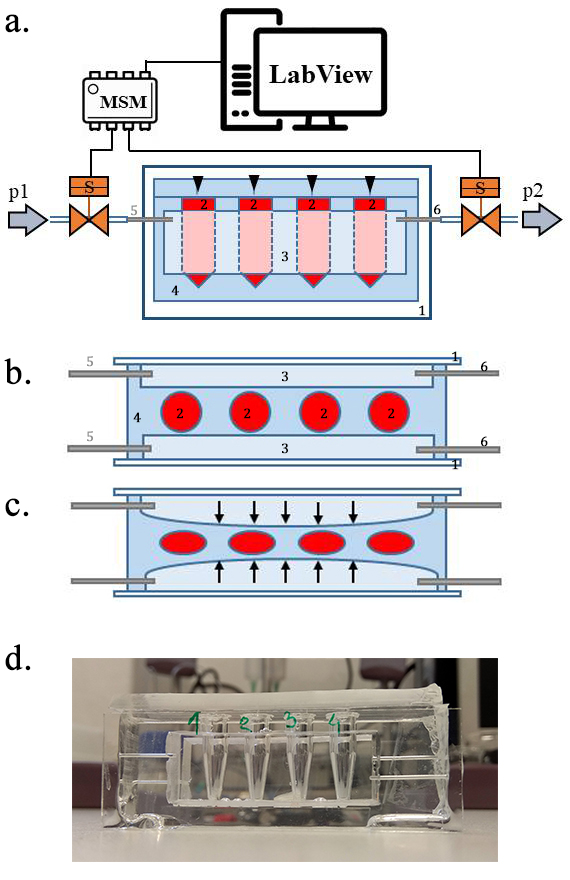

**Figure S3**. a) Scheme of the set-up with the pressure oscillator connected to the Eppendorf chip with side view and b) top view of the “Eppendorf chip”. Numbers indicating the glass slide (1), the sample chambers (2), the cuboid cavity (3), the PDMS and the inlet (5) and outlet (6) steel capillaries, respectively. b-c) Schematic representation of the chip’s operation upon applied pressure: the cavity (3) expands towards the sample chambers (2) and squeeze them. d) Photo of the constructed device prior to filling and connection to the pressure controller.

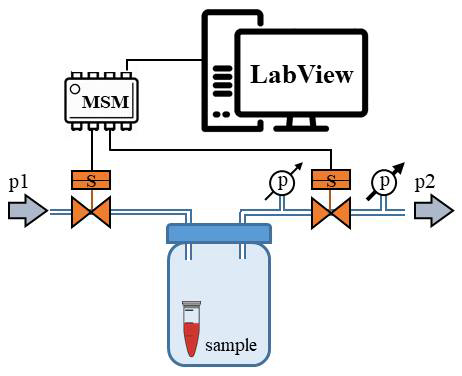

**Figure S4**. Schematic representation of the “pressure bottle” system. Two manometers were attached to the system and used to determine the pressure acting on the samples.

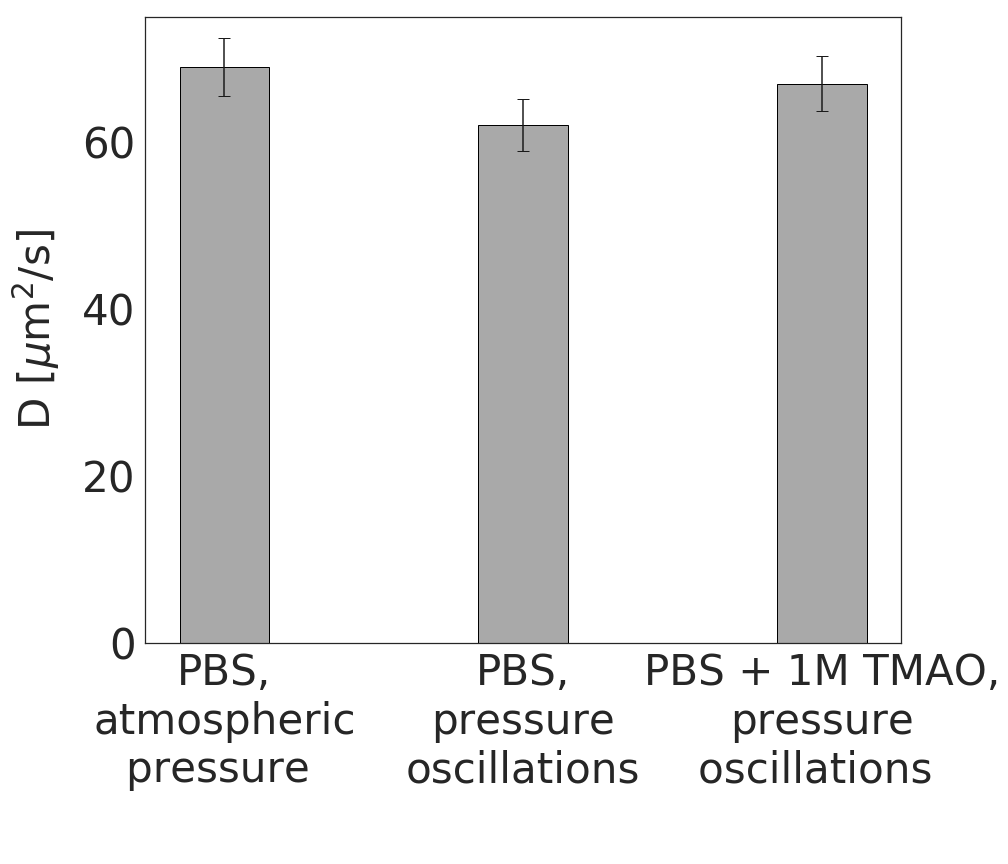

**Figure S5.** Diffusion coefficients,
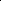
 of LDH measured by FCS. First bar is a control; the two following correspond to samples incubated for 24 hours in the high-pressure oscillation system without and with 1M TMAO in the solution.

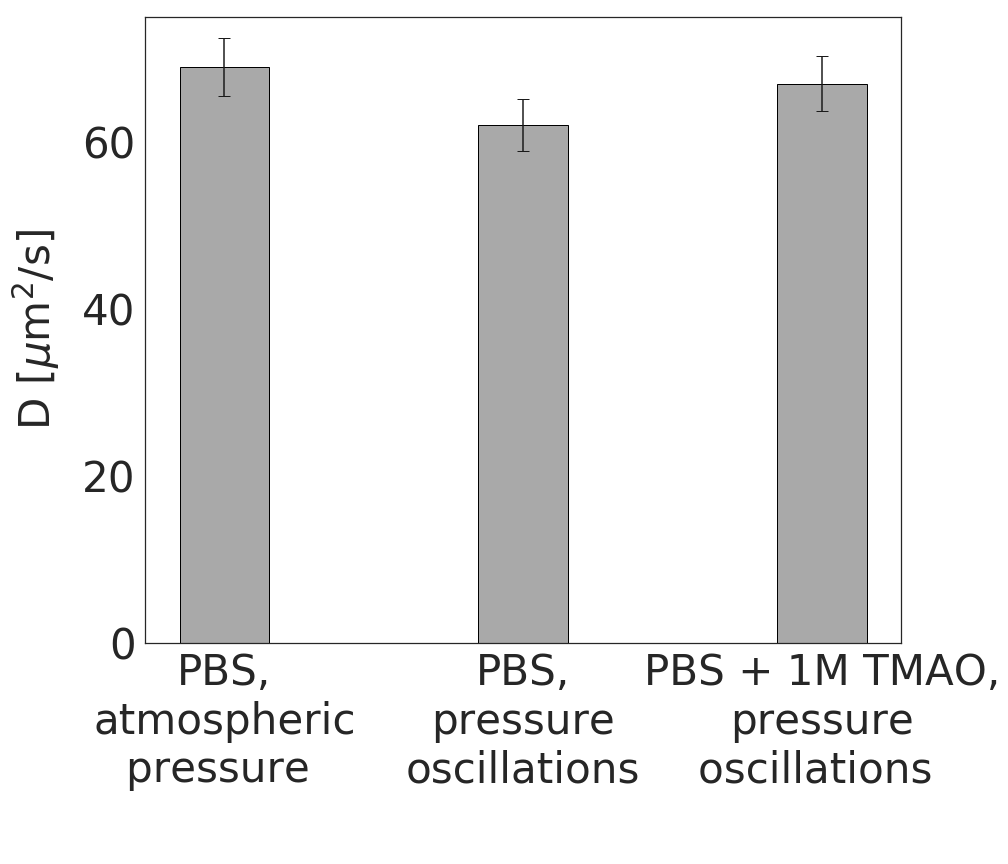

**Figure S6.** Diffusion coefficients of LDH incubated for 15 minutes at elevated temperatures, scaled by the diffusion coefficient of native (not temperature-treated) LDH. Black symbols refer to LDH incubated in pure PBS, red symbols – to LDH incubated in PBS with 1M TMAO. Symbol shapes differentiate between three independent measurement series (series 1 – squares, series 2 – circles, series 3 – triangles). Initial LDH concentration during incubation was around 3 nM in series 1 and an order of magnitude higher for series 2 and 3.

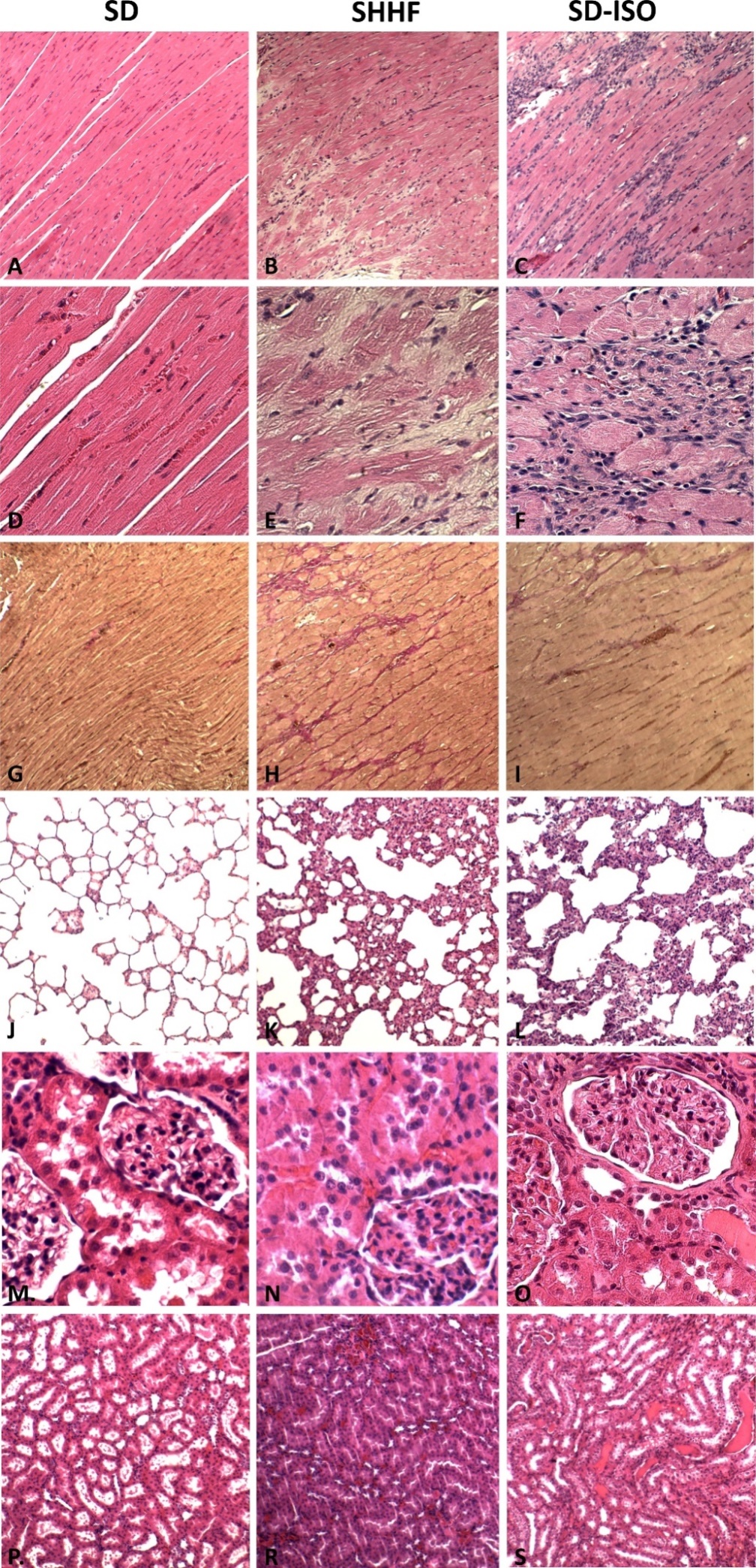

**Figure S7 Comparison of the histopathological picture between SD, SHHF and ISO-SD.**

A, B, C - myocardium; hematoxylin-eosin staining at magnification x10; D, E, F - myocardium; hematoxylin-eosin staining at magnification x40; G, H, I- myocardium; van Gieson staining at magnification x10; J, K, L - lungs; hematoxylin-eosin staining at magnificarion x10; M, N, O – kidney - renal cortex, renal bodies; hematoxylin-eosin staining at magnificarion x40; P, T, S – kidney - renal medulla; hematoxylin-eosin staining at magnification x10.
